# Multimodal deep neural decoding of visual object representation in humans

**DOI:** 10.1101/2022.04.06.487262

**Authors:** Noriya Watanabe, Kosuke Miyoshi, Koji Jimura, Daisuke Shimane, Ruedeerat Keerativittayayut, Kiyoshi Nakahara, Masaki Takeda

**Author notes:** Correspondence should be addressed to: Masaki Takeda, Ph.D. Research Center for Brain Communication, Kochi University of Technology, 185, Miyanokuchi, Tosayamada-cho, Kami-shi Kochi, 785-8502, Japan.

## Abstract

Perception and categorization of objects in a visual scene are essential to grasp the surrounding situation. However, it is unclear how neural activities in spatially distributed brain regions, especially in terms of temporal dynamics, represent visual objects. To address this issue, we explored the spatial and temporal organization of visual object representations using concurrent functional magnetic resonance imaging (fMRI) and electroencephalography (EEG), combined with neural decoding using deep neural networks (DNNs). Visualization of the fMRI DNN revealed that visual categorization (faces or non-face objects) occurred in brain-wide cortical regions, including the ventral temporal cortex. Interestingly, the EEG DNN valued the earlier phase of neural responses for categorization and the later phase of neural responses for sub-categorization. Combination of the two DNNs improved the classification performance for both categorization and sub-categorization. These deep learning-based results demonstrate a categorization principle in which visual objects are represented in a spatially organized and coarse-to-fine manner.

## Introduction

Visual perception is a rapid cognitive process that allows immediate judgement of safety/risk, enabling to quick planning and acting. The recognition and categorization of a visual scene require a series of processing stages in the ventral visual processing stream (Grill-Spector and Weiner, 2014; Grill-Spector et al., 2017; Hung et al., 2005; Logothetis and Sheinberg, 1996; Peelen and Downing, 2017). The last stage of the ventral stream, where perceptual and mnemonic signals meet, plays a crucial role in this cognitive process by associating the perceived object with the neural representations of memorized objects stored in the long-term memory (Landi et al., 2021; Suzuki and Naya, 2014; Takeda et al., 2018).

The functional role of the ventral stream on visual categorization has been investigated in neuropsychological studies. Lesions in the ventral stream can cause various forms of agnosia depending on the location of the core lesion (Barton, 2011; Cavina-Pratesi et al., 2010; Gainotti and Marra, 2011; Konen et al., 2011; Schiltz et al., 2006), suggesting module-like spatial organization for visual categories, such as faces and objects. These observations are further supported by accumulating evidence from functional magnetic resonance imaging (fMRI) in humans, such as the identification of category-selective regions for faces (Haxby et al., 2001; Ishai et al., 1999; Kanwisher et al., 1997), objects (Haxby *et al*., 2001; Ishai *et al*., 1999; Kourtzi and Kanwisher, 2000; Malach et al., 1995), and places (Epstein and Kanwisher, 1998) in not only the ventral stream, but also other cortical regions. Spatial organization in the categorization was also demonstrated in non-human primates (Kriegeskorte et al., 2008; Tsao et al., 2003; Tsao et al., 2006; Turesson et al., 2012).

In contrast, neural representations for categorical subdivision (sub-categorical representation), especially in terms of the temporal aspects, remain unclear. At the finest spatial scale, single neurons show selective responses to identities within a single category in non-human primates (Miyashita and Chang, 1988; Tanaka, 1996). At a larger spatial scale, a high-frequency local field potential carried spike-coupled information, such as the species, view, and identity of the face (Miyakawa et al., 2018). In human studies, extensive evidence has been obtained from magneto-/electroencephalography (MEG/EEG) (Carlson et al., 2013; Cichy et al., 2014; Dobs et al., 2019; Liu et al., 2002; Zheng et al., 2012) (see also fMRI studies for sub-categorical representation (Kriegeskorte et al., 2007; Nestor et al., 2011)). However, the temporal trajectories of categorical and sub-categorical representation, that is, whether our brains perform visual object categorization in a fine-to-coarse or coarse-to-fine manner, remains controversial (Carlson *et al*., 2013; Cichy *et al*., 2014; Dobs *et al*., 2019; Liu *et al*., 2002; Mack and Palmeri, 2011).

Unraveling the complete picture of the spatiotemporal profile of categorical and sub-categorical representations of visual objects is indispensable for understanding the computational model of visual object processing in the brain. To address this issue, we introduced two technical implementations in this study. First, we employed concurrent fMRI and EEG recordings in humans to obtain neural responses to visual stimuli at high spatiotemporal resolution. Second, we utilized a deep neural network (DNN) to resolve the spatiotemporal patterns of the neural activity responsible for the categorical/sub-categorical classification of visual objects. We hypothesized that (1) fMRI data involve spatially resolved patterns of neural activity that contribute to visual object classification between categories (faces or objects) and (2) EEG data involve temporally resolved patterns of neural activity that contribute to visual object classification not only between categories, but also within categories (sub-categorization; male faces or female faces, and natural objects or artificial objects), both of which would be processed at different periods (Figure 1A). These hypotheses were tested using newly developed DNN classifiers for multimodal neural data recorded by concurrent fMRI and EEG (Figure 2) as well as univariate analysis for the fMRI and EEG data.

**Figure 1.**
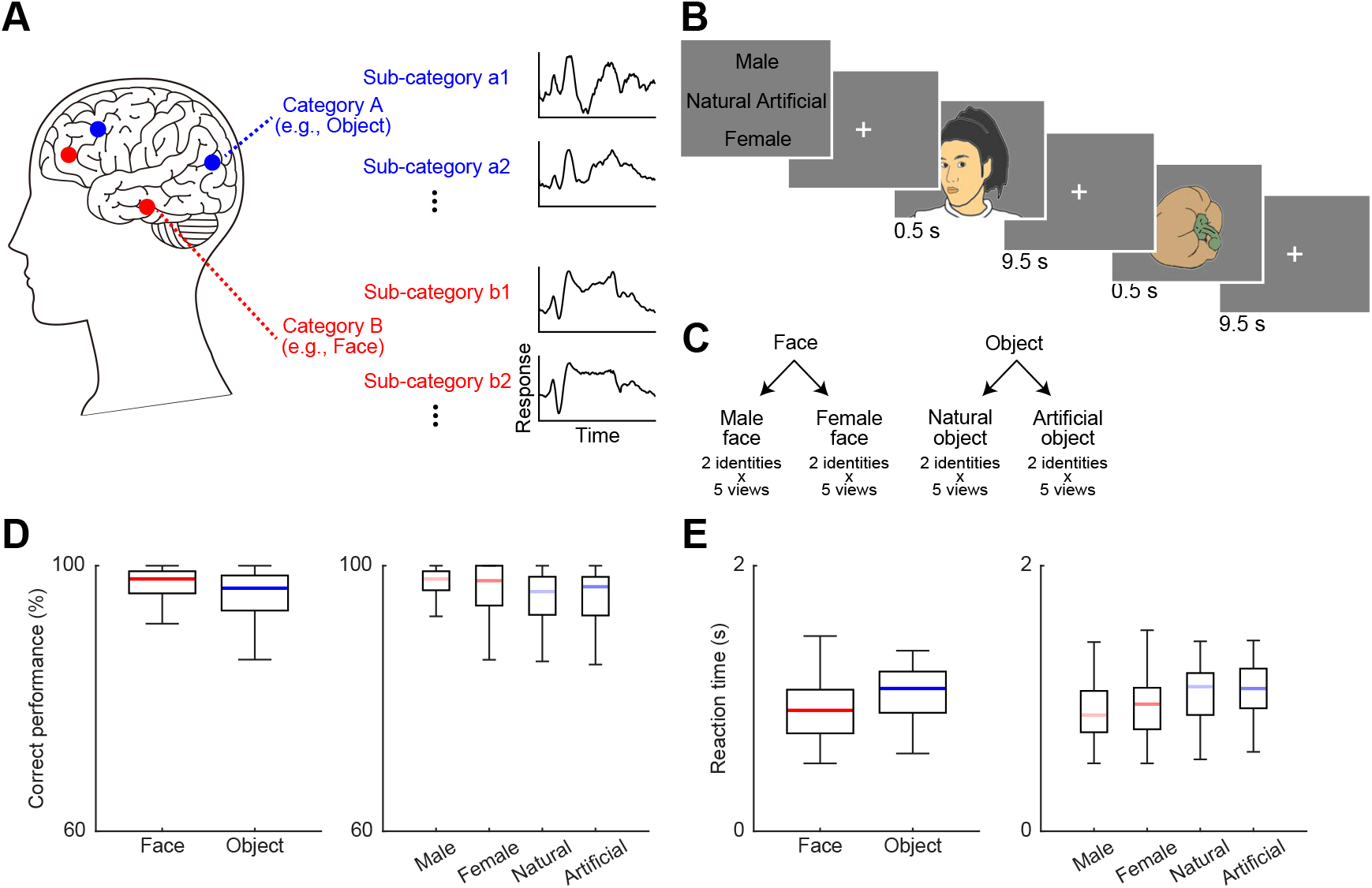
Experimental paradigm and behavioral performance. **(A)** Hypothesis for the neural representation of visual object categorization and sub-categorization. Categorical information, such as identification of faces or objects, is mainly represented in the spatial domain, whereas sub-categorical information, such as male faces or female faces is mainly represented in the temporal domain. **(B)** Sequence of a task run. At the start of a task run, instruction are given regarding the associations between the visual objects and buttons. Each visual stimulus from the four sub-categories (male face, female face, natural object, artificial object stimuli) is presented for 500 ms, with a 10-s interval. **(C)** Example visual stimuli. Face and object images are selected from ATR Facial Expression Database DB99 and ALOI, respectively. See also Figure 1 – figure supplement 1 for their low-level visual features. **(D–E)** Correct performance (D) and response latency (E) for each category and sub-category. **(D)** Differences in the correct performance in both face sub-category and object sub-category are not statistically significant (right, Wilcoxon signed rank test, *z* < 0.4, *P* > 0.68), although the performance in the face trial is higher than that in the object trial (left, face and object; *z* = 4.1, *P* = 4.19×10^-5^). **(E)** Left, the reaction times to the face stimuli are significantly faster than those to the object stimuli (Wilcoxon signed rank test, *z* = 5.56, *P* = 2.63×10^-8^). Right, within the face sub-category, the reaction times to the male face are faster than those to the female face (*z* = 3.50, *P* = 4.63×10^-4^), whereas the difference in reaction times within the object sub-category is not significant (*z* = 0.71, *P* = 0.48). *N* = 50 participants. Boxplots show median (thick line), 25^th^/75^th^ percentile (box), and minimum/maximum (whisker).

**Figure 2.**
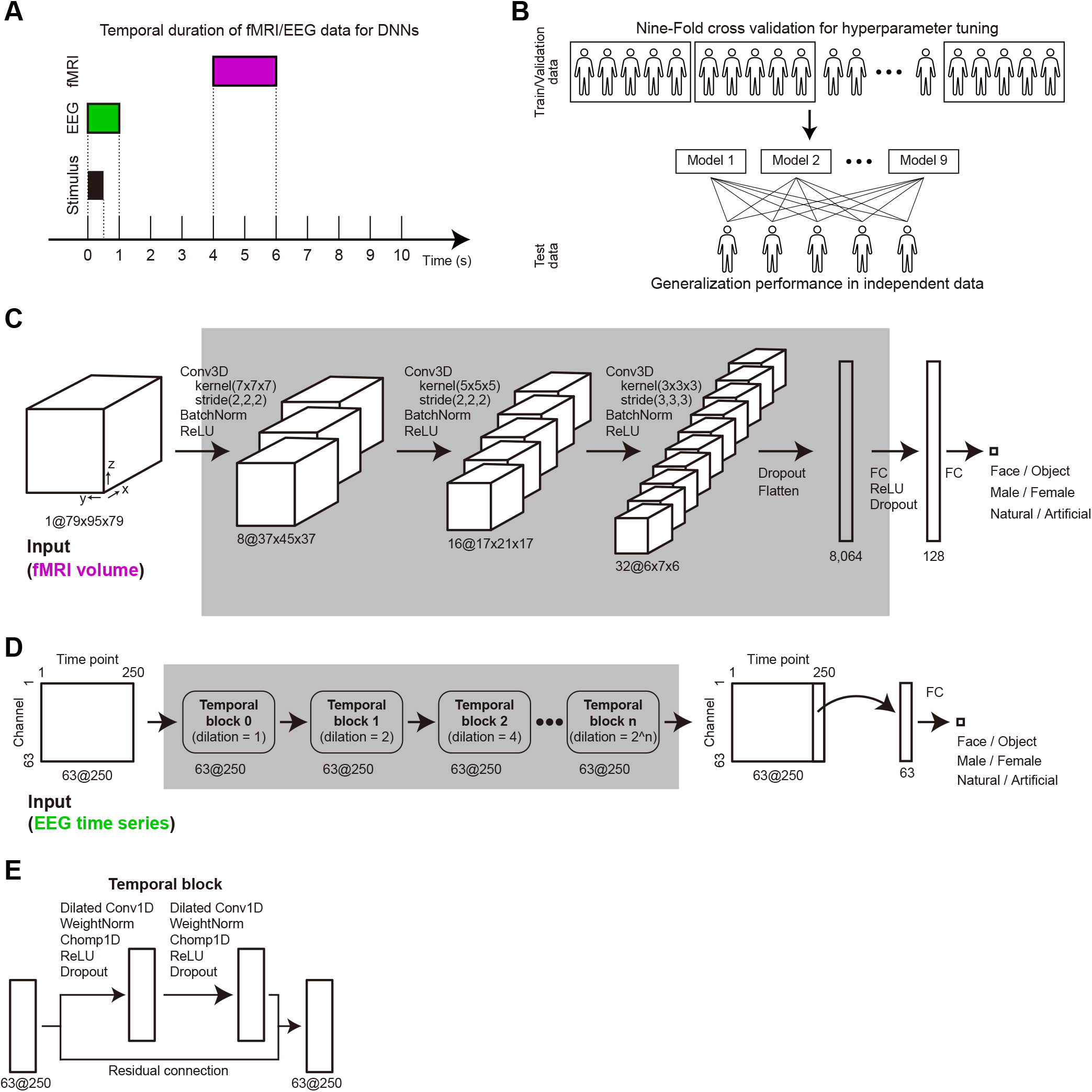
DNN models for multimodal neural data. **(A)** Temporal relationship across stimulus presentation and fMRI/EEG data windows used for DNN. The fMRI data were taken within 4–6 s from stimulus onset to consider the delayed hemodynamic response of the BOLD signal. The EEG data were taken during the period from stimulus onset to 0.5 s after stimulus offset. **(B)** Schematics of the training/validation/test architecture. In the training/validation dataset (*N* = 45 participants, one-fold data include data from *N* = 5 participants), we applied nine-fold cross-validation to tune the hyperparameters and constructed learned DNN models. Then, the generalization performances of these models for the independent test dataset were assessed (*N* = 5 participants). **(C)** DNN model for fMRI data. The model includes three convolution layers and two fully connected layers. **(D)** DNN model for EEG data. The model includes four temporal blocks and one fully connected layer. **(E)** Temporal block of the EEG DNN. This block includes two dilated convolution layers with a residual connection.

## Results

### Behavioral results

Participants performed a visual object classification task in which they judged whether the presented visual stimuli belonged to the male face, female face, natural object, or artificial object image category (Figure 1B–C). To minimize the effects of low-level visual features of stimuli on differential brain activations and resulting differences in decoding performance, we controlled for the image contrast and luminance (Stigliani et al., 2015) (Figure 1 – figure supplement 1; see Materials and Methods for details).

The participants almost perfectly categorized the presented visual stimuli (Figure 1D; *N* = 50). The performance was not significantly different between the male and female face trials or between the natural and artificial object trials (Wilcoxon signed rank test, *z* < 0.4, *P* > 0.68). Although the reaction times to male faces were faster than those to female faces (*z* = 3.50, *P* = 4.63×10^-4^), there was no significant difference in reaction times within the object sub-category (*z* = 0.71, *P* = 0.48) (Figure 1E).

### Spatial organization of category-specific information revealed by fMRI activations

We explored brain regions associated with categorical processing based on standard univariate general linear model (GLM) analysis of fMRI data (Figure 3, Figure 3 – figure supplement 1). Robust activations for faces compared with those for objects were observed in various regions, including the temporal occipital fusiform cortex (TOF), which includes the fusiform face area (FFA), and lateral occipital complex (LOC) in the ventral temporal cortex (VTC), as well as the occipital pole, angular gyrus, frontal medial cortex, amygdala, frontal orbital cortex, precuneous (PcC), and middle temporal gyrus (Figure 3A). In contrast, the brain regions activated by objects included the TOF, LOC, middle frontal gyrus, insula, superior frontal gyrus, thalamus, and paracingulate gyrus. These results were consistent with those of previous studies (Haxby *et al*., 2001; Ishai *et al*., 1999; Kanwisher *et al*., 1997; Kourtzi and Kanwisher, 2000; Malach *et al*., 1995).

**Figure 3.**
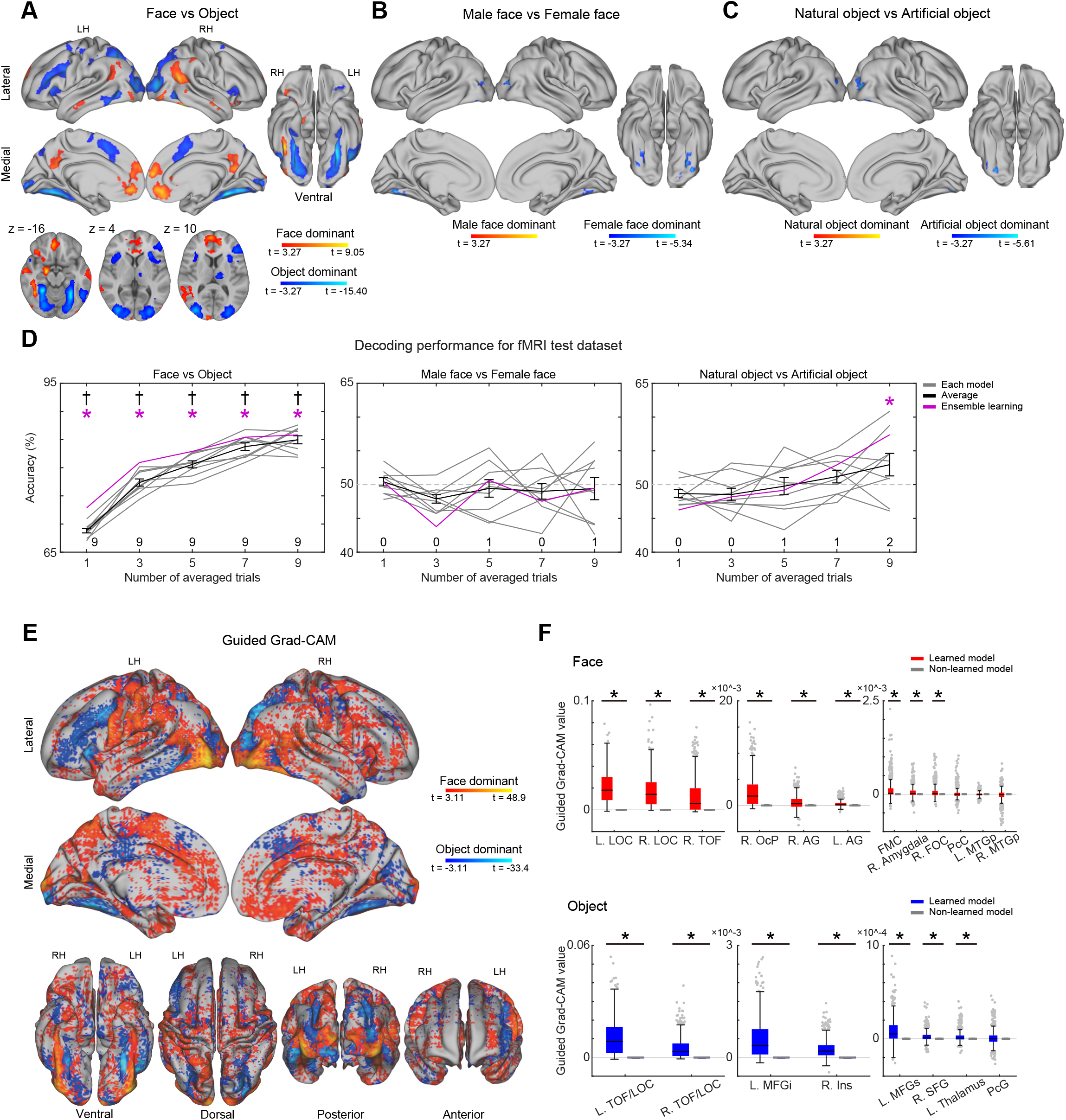
Spatial representation of categorization/sub-categorization for visual objects revealed by fMRI. **(A–C)** Activation maps for the contrast between the face and object (A), male face and female face (B) and natural object and artificial object (C) trials. The color bars denote t-values. The significant brain regions were identified using a threshold of *P* < 0.05 corrected with a cluster-wise family-wise error rate based on a non-parametric permutation test. The identified clusters in the subcortical structures and the insula are shown in the horizontal sections. See also Figure 3 – figure supplement 1–3. **(D)** Generalization decoding performance for the test dataset. The ensemble accuracy (purple) as well as average accuracy ± S.D (thick line) across models are shown as a function of the number of averaged trials. The thin lines represent the accuracy in each model. *, binomial test for ensemble accuracy against chance level (50%), FDR corrected *q* < 0.05. †, t-test for accuracy distribution against chance level, FDR corrected *q* < 0.05. The number of significant models (binomial test, *q* < 0.05) is shown above the abscissa. **(E)** Spatial organization of the guided Grad-CAM values for the contrast of category (between face and object trials). The threshold of significance was set at *P* < 0.05 corrected with cluster-wise FWE based on a non-parametric permutation test. **(F)** ROI analysis. Colored and grayed bars denote the guided Grad-CAM values in learned and non-learned models, respectively. For display purposes, the results are plotted in separate panels according to the magnitudes of the values. *, t-test against values obtained from the non-learned model, FDR corrected *q* < 0.001. Values in non-learned models are nearly zeros. Boxplots show median (black line), 25^th^/75^th^ percentile (box), nonoutlier minimum/maximum (whisker), and outlier (dot, data points outside 1.5× interquartile range). The cluster information is listed in Figure 3 – figure supplement 1. L, left; R, right; LH, left hemisphere; RH, right hemisphere; AG, angular gyrus; FMC, frontal medial cortex; FOC, frontal orbital cortex; Ins, insular cortex; LOC, lateral occipital cortex; MFG, middle frontal gyrus; MTG, middle temporal gyrus; OcP, occipital pole; PcC, precuneous cortex; PcG, paracingulate gyrus; SFG, superior frontal gyrus; TOF, temporal occipital fusiform cortex; i, inferior; s, superior; p, posterior.

Subsequently, we explored brain regions associated with sub-categorical processing (Figure 3 – figure supplement 2, 3). The face gender contrast (male face vs. female face) revealed weak activations; there were no male face-dominant clusters and three female face-dominant clusters (Figure 3B). The contrast between natural and artificial objects also showed weak activations; there were no natural object-dominant clusters and two artificial object-dominant clusters (Figure 3C). These results suggest that the differences in the sub-categorical processing of visual objects are hardly detected at the spatiotemporal resolution of fMRI used in this study.

### Deep neural decoding of visual object categorization by fMRI DNN

To decode information about the visual object category and identify the spatial organization of the responsible brain regions, we used a DNN classifier for the fMRI data, which was based on three-dimensional convolution layers (Figure 2C). We trained the fMRI DNN using an fMRI volume acquired 4 s after the presentation of visual stimuli in each trial (Figure 2A). Training and validation of the DNN were conducted by nine-fold cross-validation, and the generalization performance of the resultant nine models was evaluated using an independent test dataset (Figure 2B; see Materials and Methods for details).

The ensemble learning method using a majority voting scheme (Rokach, 2009) revealed that the classification performance in categorization reached statistical significance when the trials were not averaged (72.90%; *q* = 2.61×10^-72^, binomial test against chance level 50%, corrected for multiple comparisons by the false discovery rate (FDR) of *q* < 0.05) (Figure 3D). The performance improved as a function of the number of averaged trials, reaching 85.83% with an average of nine trials (*q* = 3.89×10^-185^). In contrast, neither face nor object sub-categorical classification was significant in the averaged trial condition (*q* > 0.21), except for object sub-categorization when nine trials were averaged.

Two additional analyses were performed to examine the robustness of these results. In the first analysis, a t-test showed that for categorical classification, the performance distribution statistically reached above chance when trials were not averaged [68.80 ± 0.39% (mean ± s.e.m.); *t* = 48.80, *P* = 3.44×10^-11^, corrected by FDR *q* < 0.05]. The performance improved as a function of the number of averaged trials (linear regression, *R*^2^ = 0.82, slope coefficient, *t* = 13.99, *P* = 1.32×10^-17^), reaching 84.96 ± 0.71% with an average of nine trials (*t* = 49.04, *P* = 3.31×10^-11^). The highest accuracy reached was 87.60% in the best performing DNN model. In contrast, neither face nor object sub-categorical classification reached statistical significance in either averaged set of trial conditions (*q* > 0.26).

In the second analysis, we counted the number of models showing significant performance against chance level using a binomial test (threshold at *P* = 0.05, FDR corrected). Although all nine models were significant in categorical classification across all averaged trial conditions, less than two models had significant sub-categorical classification. The results of the two additional analyses support those of the ensemble learning method.

With respect to important spatial neural activity for visual object classification, guided gradient-weighted class activation mapping (Selvaraju et al., 2019) (guided Grad-CAM) showed that the fMRI DNN model valued brain-wide regions, including the frontal, parietal, temporal, and occipital cortices as well as subcortical structures in both the face and object trials (Figure 3E; Figure 3 – figure supplement 4). The whole-brain activations identified in the univariate analysis and guided Grad-CAM were significantly correlated at the voxel-by-voxel level, although most voxels showed guided Grad-CAM values near zero, leading to a relatively low effect size (Figure 3 – figure supplement 5A; Spearman’s rank correlation, *ρ* = 0.048, *t* = 19.73, *P* < 0.001 for face trial; *ρ* = −0.21, *t* = −90.13, *P* < 0.001 for object trial). The correlation coefficient increased when the target voxels were masked by the meta-analysis maps obtained from Neurosynth (Yarkoni et al., 2011) using the terms “face” and “object” (Figure 3 – figure supplement 5B; *ρ* = 0.27, *t* = 46.27, *P* < 0.001 for face trial; *ρ* = −0.30, *t* = −56.21, *P* < 0.001 for object trial). This finding suggests that voxels with nearly zero values in the guided Grad-CAM tended to be located outside significant clusters in the univariate analysis. Indeed, the region-of-interest (ROI) analysis revealed that most of the clusters identified in the univariate analysis had significant guided Grad-CAM values in both the face and object trials compared with those obtained in non-learned DNN models in which weight values were assigned randomly (Figure 3F; Wilcoxon signed rank test against guided Grad-CAM values in non-learned models, FDR corrected *q* < 0.05). Similar results were obtained when ROIs were defined based on the anatomical structure or functional meta-analysis by Neurosynth (Figure 3 – figure supplement 6); among the three ROI-based analyses, consistent results were obtained in the VTC (LOC, TOF), FMC, and PcC in the face trial, and the VTC (LOC), and thalamus in the object trial.

These results indicate that, to categorize faces and objects, our DNN model paid attention to the neural signals in the distributed brain regions identified in the univariate analysis.

### Temporal trajectory of categorical/sub-categorical information by EEG responses

To examine the temporal characteristics of categorical and sub-categorical processing of visual objects, we analyzed the event-related potential (ERP) and power spectrogram of the EEG signals (Figure 4; Figure 4 – figure supplement 1). Regarding the ERPs for categorization, the face stimuli induced an early negative component (< 200 ms from stimulus onset) with a prolonged component (approximately 200–700 ms), wherein the polarity of the potential and its dynamics differed across channels. In contrast, the object stimuli did not evoke a clear early negative component (Figure 4 – figure supplement 1A). This characteristic led to differential ERPs between the face and object trials (Figure 4A). The ERPs between the faces and objects were significantly different at approximately 200–600 ms from stimulus onset in the brain-wide (50/63) channels (Figure 4 – figure supplement 1B; cluster-defining threshold of *P* < 0.05, corrected significance level of *P* < 0.05). In contrast, the power spectrogram did not show clear differences (Figure 4A; Figure 4 – figure supplement 1B).

**Figure 4.**
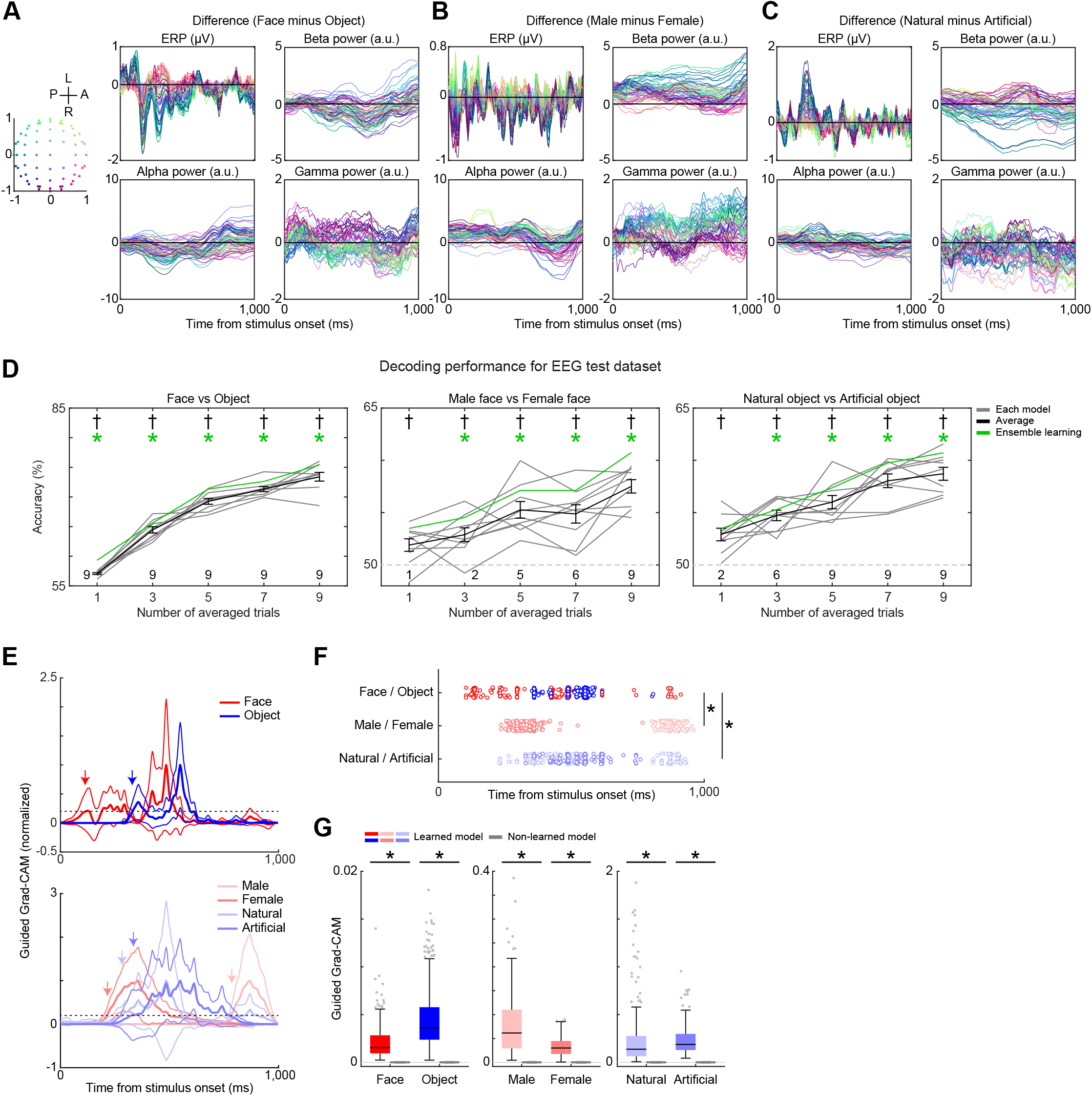
Temporal representation of categorization/sub-categorization for visual objects revealed by EEG. **(A)** Differential ERP and power spectrogram in the contrast of face minus object trials. The line colors correspond to the channel locations depicted on the left side. A, anterior, P, posterior, R, right, L, left. **(B)** Differential ERP and power spectrogram in the contrast of male face minus female face trials. **(C)** Differential ERP and power spectrogram in the contrast of natural object minus artificial object trials. **(D)** Generalization decoding performance for the test dataset. The configurations are the same as those in Figure 3d. **(E)** Temporal trajectories of the guided Grad-CAM values for each trial type. The data from all channels were averaged, and values were normalized to the maximum guided Grad-CAM value (=1). The dashed lines represent the threshold (20% of the maximum value). The arrows depict the maximum latency that first exceeded the threshold. **(F)** Temporal distribution of maximum latency for each trial (dot). The trials for the face, object, male face, female face, natural object, and artificial object are separately colored. *, Tukey–Kramer test, *P* < 0.05 after Kruskal–Wallis test, *χ*^2^ = 9.70, *P* = 0.0078. **(G)** Comparison of the guided Grad-CAM values at the maximum latency between the learned and non-learned models. *, t-test against values obtained from the non-learned model, FDR corrected *q* < 0.001. Note that the values in non-learned models are nearly zero. The boxplots show the median (black line), 25^th^/75^th^ percentile (box), nonoutlier minimum/maximum (whisker), and outlier (dot, data points outside 1.5× interquartile range).

In contrast to categorization, the spectral power was more informative in sub-categorization. Specifically, in face sub-category processing, the male face stimuli elicited greater beta power than the female face stimuli in a prolonged time window (– 1,000 ms). However, these two stimuli did not evoke distinct ERP responses (Figure 4B and Figure 4 – figure supplement 1C). In object sub-category processing, artificial stimuli evoked larger beta power than natural stimuli in a prolonged time window (– 1,000 ms) in several channels at the posterior part of the brain. Distinctive ERP responses were also observed at 200–300 ms (21/63 channels; Figure 4D and Figure 4 – figure supplement 1D).

### Deep neural decoding of visual object categorization/sub-categorization by EEG DNN

To decode information about the visual object category and sub-category as well as to identify its temporal dynamics, we used a DNN classifier for the EEG data, which was based on a temporal convolutional network (TCN) (Bai et al., 2018) (Figure 2D–E). We trained the EEG DNN using a 63-ch EEG time series during 1 s after stimulus onset (Figure 2A). The ensemble learning method revealed that the classification performance in categorization was statistically above the chance when the trials were not averaged (59.33%; binomial test against chance level 50%, *q* = 5.75×10^-13^, corrected for multiple comparisons by FDR *q* < 0.05) (Figure 4D). The performance improved as a function of the number of averaged trials, reaching 75.41% with an average of nine trials (*q* = 4.95×10^-89^). Regarding face and object sub-categorization, the classification performance improved from 53.50% (no trial average; *q* = 0.056) to 60.75% (average of nine trials; *q* = 1.25×10^-8^) and from 53.48% (no trial average; *q* = 0.067) to 60.72% (average of nine trials; *q* = 4.99×10^-8^), respectively. These results suggest that the EEG DNN succeeded in decoding the sub-categorization of visual objects whereas the fMRI DNN did not.

The performance distribution and number of significant models were further analyzed to confirm the robustness of the results. First, the performance distributions of nine models for categorization statistically reached above chance when the trials were not averaged (57.07 ± 0.15%; t-test against chance level 50%, *t* = 47.33, *P* = 4.39×10^-11^, FDR corrected *q* < 0.05) (Figure 4D). The performance improved as a function of the number of averaged trials (linear regression, *R*^2^ = 0.87, slope coefficient, *t* = 16.90, *P* = 1.34×10^-20^), resulting in 73.41 ± 0.74% with an average of nine trials (*t* = 31.73, *P* = 1.06×10^-9^). The decoding performance for both face and object sub-categorization also improved as the number of averaged trials increased (face sub-categorization: *R*^2^ = 0.44, slope coefficient, *t* = 5.78, *P* = 7.51×10^-7^; object sub-categorization: *R*^2^ = 0.59, slope coefficient, *t* = 8.00, *P* = 4.78×10^-10^). Regarding male face/female face classification, the mean accuracy under the condition in which the trials were not averaged was significantly above chance (51.90 ± 0.60%; *t* = 3.19, *P* = 0.0129, FDR corrected). When nine trials were averaged, the accuracy reached 57.51 ± 0.64% (*t* = 11.76, *P* = 2.49×10^-6^). Notably, the highest accuracy was 59.46% in the best performing model. Regarding natural object/artificial object classification, the accuracy improved from 52.91 ± 0.60% (no trial average; *t* = 4.86, *P* = 0.0013, FDR corrected) to 58.71 ± 0.62% (average of nine trials, *t* = 14.08, *P* = 6.30×10^-7^). The highest accuracy was 61.56% in the best performing model.

Second, while all nine models were significant in categorical classification across all averaged trial conditions, the number of models with significant sub-categorical classification increased as a function of the number of averaged trials (binomial test, threshold at *P* = 0.05, FDR corrected). In face sub-categorization, the number of significant models increased from one to nine, whereas in object sub-categorization, the number increased from two to nine.

Visualization by guided Grad-CAM revealed that our DNN model valued differential periods between categorization and sub-categorization (Figure 4E). The model started to value the timepoints at 120 ms and 336 ms in the face and object trials, respectively (“emergence latency”; see Materials and Methods for details). Regarding sub-categorization, the four emergence latencies were 788 ms for male face trials, 216 ms for female face trials, 288 ms for natural object trials, and 328 ms for artificial object trials, all of which were slower than that for face trials. Meanwhile, emergence latencies for female face and natural object trials were longer than those for object trials. This “categorization-then-subcategorization” tendency in the latency of first valued timepoints was also observed in the latency with the maximum guided Grad-CAM value (“maximum latency”) (Figure 4F). The maximum latency varied among the trials, and their ranges differed between categorization and sub-categorization: the maximum latency in categorization varied between 104 ms and 912 ms (median = 498 ms), whereas those in face and object sub-categorization varied between 236 ms and 960 ms (median = 556 ms) and between 228 ms and 932 ms (median = 541 ms), respectively. The maximum latency in categorization was significantly shorter than those in face and object sub-categorization (Kruskal–Wallis test, *χ*^2^ = 9.70, *P* = 0.0078; post-hoc Tukey– Kramer test, *P* < 0.05). Notably, the maximum guided Grad-CAM values in each trial type were significantly greater than those in the non-learned models (Figure 4G; Wilcoxon signed rank test against guided Grad-CAM values in non-learned models, FDR corrected *q* < 0.05). When grouping the guided Grad-CAM values into six brain regions based on electrode channel locations, all regions showed significant guided Grad-CAM values (Figure 4 – figure supplement 2). Notably, the highest guided Grad-CAM value in categorization was for the occipital cortex, whereas the frontal and parietal cortices were more valued in sub-categorization (Friedman’s test with post-hoc Tukey-Kramer method, *P* < 0.05).

Collectively, these results indicate that our EEG DNN model successfully captured not only the categorical representation, but also the sub-categorical representation of visual objects in the temporal domain of neural responses. Our results experimentally support that visual object categorization be performed in a coarse-to-fine manner along the temporal axis.

### Concurrent neural decoding by combined DNN model

Simultaneous acquisition of fMRI and EEG data enables concurrent neural decoding using an integrated DNN. Thus, we performed neural decoding for concurrent fMRI and EEG data with the combined fMRI–EEG DNN (Figure 5A). In the DNN, feature extraction layers, three-dimensional (3D) convolution layers for fMRI data and temporal blocks for EEG data were finally concatenated and fully-connected.

**Figure 5.**
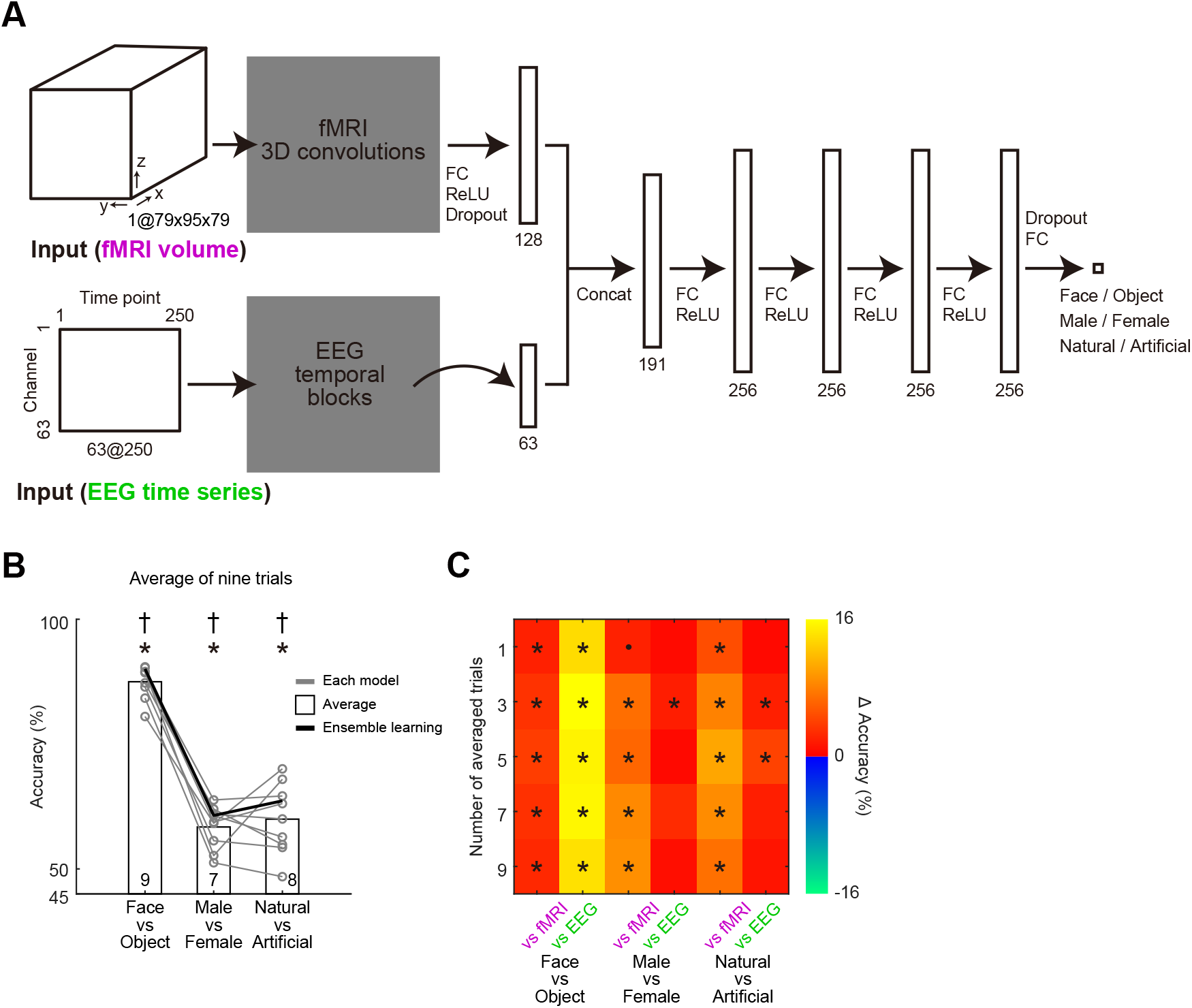
Combined fMRI–EEG DNN model. **(A)** Architecture of the model. Firstly, the fMRI volume data and EEG time series data are separately convolved by the conv3Ds and temporal blocks shown in Figure 2. Next, the resulting outputs are concatenated, are then processed using four fully connected layers. **(B)** Generalization decoding performance for the test dataset (average of nine trials). The bars denote the mean accuracy across models, and the circles denote the accuracy of each model. *, binomial test for ensemble accuracy against chance level (50%), FDR corrected *q* < 0.05. †, t-test for accuracy distribution against chance level, FDR corrected *q* < 0.05. The number of significant models (binomial test, *q* < 0.05) is shown above the abscissa. **(C)** Improvement in decoding accuracy of fMRI–EEG DNN against the accuracies of the fMRI and EEG DNN models. The differences in accuracy are color-coded. Note that neither the fMRI DNN nor EEG DNN shows higher accuracy than the fMRI–EEG DNN. *, t-test, FDR corrected *q* < 0.05. *, *P* < 0.05, uncorrected.

When the fMRI and EEG activities of nine trials were averaged, the ensemble performance of the fMRI–EEG DNN was 90.05%, 60.70%, and 63.64% for categorization, face sub-categorization, and object sub-categorization, respectively, all of which were significantly above the chance level (Figure 5B; binomial test, *P* = 3.03×10^-240^, *P* = 2.51×10^-9^, and *P* = 2.44×10^-13^). The performance distributions in the nine models were also significant against the chance level (categorical, 87.49 ± 1.10%, *t* = 78.97, *P* = 7.37×10^-13^; face sub-categorical, 58.40 ± 1.43%, *t* = 40.84, *P* = 1.42×10^-10^; object sub-categorical, 59.97 ± 2.36%, *t* = 25.44, *P* = 6.11×10^-9^; all *q* < 0.05 in FDR correction). The best performing models showed performances of 90.46%, 63.81%, and 70.04%. A binomial test revealed that nine, seven, and eight models achieved significant decoding performance for categorization, face sub-categorization, and object sub-categorization, respectively (threshold at *P* = 0.05, FDR corrected).

In the comparison of the classification accuracy of the combined fMRI–EEG DNN with the accuracies with fMRI data alone (Figure 3) or EEG data alone (Figure 4), the fMRI–EEG DNN accuracy was higher than those on the fMRI DNN and EEG DNN in all averaged trial conditions (Figure 5C). The categorical classification accuracy of the fMRI–EEG DNN was significantly increased in all averaged trial conditions (t-test, FDR corrected *q* < 0.05). Regarding sub-categorical classification, the fMRI–EEG DNN showed significantly higher performance compared to the fMRI DNN in all averaged trial conditions (*q* < 0.05). However, the difference in performance between the fMRI– EEG DNN and EEG DNN did not reach statistical significance except for when three trials were averaged. These results suggest that concurrent usage of spatial and temporal neural data contributes to improving the neural decoding of visual objects.

## Discussion

This study revealed that the categorical representation of visual objects is supported by spatially organized brain-wide activations in a coarse-to-fine manner. To our knowledge, the present study is the first to examine neural representation of visual objects using multimodal DNNs. This approach enabled highly resolved brain-wide spatiotemporal neural decoding. The present findings will contribute to the construction of a novel paradigm for trial-based neural decoding of internal processes in patients with impaired consciousness in clinical practice.

DNNs, which have achieved state-of-the-art results for the large-scale image recognition (Simonyan and Zisserman, 2015), are now being applied to neural data, in what is known as neural decoding (Hong et al., 2016; Horikawa and Kamitani, 2017; Yamins and DiCarlo, 2016). In an fMRI DNN, we managed to use 3D voxel data as inputs, whereas some previous studies used 2D slice data (Sarraf et al., 2017) or 2D flattened cortical data as inputs (Tsumura et al., 2021b) (see also other studies in which 3D voxel data were used as input data (Frey et al., 2021; Vu et al., 2020; Wang et al., 2020; Zhao et al., 2018)). Such image-like input could be learned by DNNs that have already been trained with a large-scale image set (e.g., ImageNet), such as VGG. The re-training approaches such as fine-tuning and transfer learning help improve the decoding performance. Given that trained models for spatio-temporal data, which are suitable for application to concurrent fMRI and EEG data, are currently unavailable, we could not employ such a strategy in this study. However, there are several clear advantages of our approach, such as (1) inclusion of neural activities in the subcortical structures, which are ignored in the 2D flattened map and (2) coverage of whole brain structures, which is impossible in 2D slice data. Indeed, we could identify the involvement of the amygdala and thalamus in the categorical classification of visual objects. Accumulating open 3D-voxel data available for deep learning is a promising means of improving 3D fMRI decoding using DNNs further.

In the EEG DNN, we employed a TCN (Bai *et al*., 2018) in which time-series data were convoluted in a dilated manner. The TCN had excellent performance for non-neural data such as character-level and word-level language modeling. Successful application of the TCN to EEG data in the present study could expand the possibility for the application of TCNs to other biological data such as electromyography data.

We obtained high decoding performance for both categorical and sub-categorical classification (*c.f.,* in the fMRI–EEG DNN, the ensemble performance reached 90.05%, 60.70%, and 63.64% for the categorical and two sub-categorical classifications, and the best performing model yielded 90.46%, 63.81%, and 70.04%). It is difficult to directly compare the performance in this study with that in other studies because the experimental paradigms (e.g., decoding technique, stimulus, and sample size) were quite different. Nevertheless, the present finding that the fMRI–EEG DNN outperforms the fMRI DNN and the EEG DNN indicates that a larger number of modal data would improve the precision of identification of the neural representation of visual objects.

In machine learning including deep learning, the learning performance is usually evaluated once for the test dataset (Raschka, 2020). In this study, we employed the ensemble learning approach (Rokach, 2009) in which obtained classification labels from nine models were voted and the majority label was treated as the predicted label for the trial. We confirmed the robustness of this ensemble approach using two distinct indices, the performance distribution and number of significant models. Considering the inherent properties of neural data to be noisy and variable between trials, this combinational approach is suitable for the application of deep learning to neural data. For example, when comparing an fMRI volume or an EEG time series with pictures in ImageNet, neural data have some background noise, c.f., it is hardly possible to detect the internal process of the subject by glimpsing an fMRI volume or an EEG time series. In addition, it is difficult to collect adequate amount of neural data that are comparable to the number of pictures in ImageNet (>10,000,000). Noisiness and a smaller sample size would result in higher variance in the decoding performance. The ensemble learning approach would reduce the variance of classification errors, and in addition to a point estimate of the classification performance, our approach can evaluate the performance variance across models.

Brain regions in which the fMRI DNN valued for categorization were closely matched with regions identified in the fMRI univariate analysis, including the ventral temporal cortex (FFA for face stimuli and LOC for object stimuli), which is consistent with previous findings (Haxby *et al*., 2001; Kanwisher *et al*., 1997; Kourtzi and Kanwisher, 2000; Malach *et al*., 1995). Notably, we revealed the involvement of the prefrontal cortex and amygdala, which have been also demonstrated in monkey (Freedman et al., 2001) and human (Kreiman et al., 2000) electrophysiology. Considering that the distribution of significant voxels is wider in the guided Grad-CAM than in the univariate analysis, deep learning techniques would have greater sensitivity to responsible brain regions for visual object categorization.

Fine and coarse information processing in visual object categorization remains controversial (Carlson *et al*., 2013; Cichy *et al*., 2014; Dobs *et al*., 2019; Liu *et al*., 2002; Mack and Palmeri, 2011). Our deep learning-based approach provided robust evidence regarding this issue: the temporal trajectory of neural decoding by the EEG DNN demonstrated that the neural activity firstly conveyed categorical information, followed by sub-categorical information. These results are in agreement with the theoretical model of “coarse-to-fine” categorization (Bar, 2004; Rousselet et al., 2002; Serre et al., 2007). One notable observation in this study is that peak time in the guided Grad-CAM value (maximum latency) varied among trials, which could be caused by the temporal variability of the behavioral states (e.g., attention) among trials. The EEG DNN could detect such variability at any time point owing to the shift invariance in the temporal domain by dilated convolutions. The trial-dependent latencies of the category/sub-category information in the neural responses would be related to the persistent time windows with significant object decoding reported in other studies (Cichy *et al*., 2014; Dobs *et al*., 2019).

Compared with the situation in which fMRI and EEG recordings are separately conducted, concurrent fMRI/EEG recording has several advantages. First, the variance of behavioral outcomes such as the correct performance and reaction time could be lower than those obtained in separate sessions or on different days. Second, concurrent usage of fMRI and EEG data can result in better decoding performance than the usage of fMRI alone or EEG alone, as shown in this study. It is promising to add other bio-signals to detect other aspects of information of visual objects such as valence or arousal.

An advanced M/EEG–fMRI fusion approach has recently attracted attention as it links multivariate neural response patterns based on representational similarity (Cichy and Oliva, 2020; Cichy *et al*., 2014), whereas M/EEG and fMRI data are acquired in separate sessions. The combination of concurrent multimodal recording and representational similarity-based fusion approach would be useful for more accurately elucidating neural representations in the sensory system.

## Materials and Methods

### Participants

A total of 53 healthy right-handed subjects (34 males and 19 females; age, 18–25 years) with normal or corrected-to-normal vision participated in the study. Of them, data from three participants were excluded due to low behavioral performance: two participants had low correct performance (< 80% at least for one sub-category) and one participant had slow responses (mean response latency > 1.5 s). The participants received 1,000 yen per hour for each session. The target number of participants was determined based on the number of trials required in previous deep learning research in neuroscience (e.g., Supratak et al (Supratak et al., 2017)) before starting the collection of neuroimaging data. This study was approved by the institutional review board of Kochi University of Technology, Kami, Japan and complied with the Declaration of Helsinki. Written informed consent was obtained from all participants.

### Experimental design

#### Stimuli

The stimuli consisted of color images (size, 7.4° × 5.8°) and a white fixation cross (size, 0.3° × 0.3°). Forty color images from face and inanimate object categories were used as the experimental stimuli. The 20 face and 20 object images were selected from ATR Facial Expression Database DB99 (http://www.atr-p.com/products/face-db.html) and Amsterdam Library of Object Images (ALOI; https://aloi.science.uva.nl/) (Geusebroek et al., 2005), respectively. The facial images were obtained from four identities: two male individuals (male face sub-category) and two female individuals (female face sub-category), with neutral facial expressions and five different angles. The object images were obtained from four identities: two natural objects (paprika and broccoli; natural object sub-category) and two artificial objects (hat and whistle; artificial object sub-category), with five different angles.

All images were contrast- and luminance-matched using in-house MATLAB (MathWorks, Natick, MA) code (Keerativittayayut et al., 2018) adapted from the SHINE toolbox (*lumMatch* function) (Willenbockel et al., 2010) for each RGB channel, after figure-ground separation. Based on a prior study (Stigliani *et al*., 2015), we evaluated (1) contrast, (2) luminance, (3) similarity to other stimuli of the same category, (4) visual field coverage, and (5) spatial frequency power distributions, after converting RGB color images into grayscale images by using the *rgb2gray* function in MATLAB (Figure 1 – figure supplement 1). In addition, hue histograms of color images were evaluated for each sub-category.

#### Task procedure

During the recording of brain activities by concurrent fMRI and EEG, the participants performed a visual object classification task, in which sequences of experimental stimuli were presented. The participants judged the corresponding visual sub-categories (male face, female face, natural object, and artificial object) and pressed a corresponding button (Figure 1B–C). At the beginning of each scanning run, a task instruction was presented that indicated the correspondences among four positions of buttons and the four sub-categories. To exclude the possibility that the learning of stimulus-response mapping would affect the neural activity, the correspondence was pseudorandomized across runs. In the scanning run, a fixation cross was firstly presented for 30 s, followed by a sequence of 50 trials. In 40 trials, face or object images were presented, and in 10 trials, blank images were presented to improve dissociation of hemodynamic responses between trials. In each trial, a stimulus was presented for 500 ms, followed by a fixation cross for 9,500 ms. Thus, each scanning run took a total of 530 s. All stimulus images were presented once in each scanning run, and the order of stimulus presentation was pseudorandomized. No visual feedback was presented. Participants practiced the task before starting concurrent fMRI and EEG.

The participants conducted 9.96 ± 2.88 runs (mean ± SD; total 496 runs) while simultaneous fMRI and EEG recordings were performed. Neural data were accordingly obtained from a total of 24,800 trials (9,920 face trials, 9,920 object trials, and 4,960 blank trials). Trials wherein the participants did not respond within 2 s (1,005 trials, 5% of total face/object trials) were excluded. The correct performance and reaction time were calculated for each category and sub-category (Figure 1D–E). Only the data from the correct trials were used for the succeeding analyses (18,007 face and object trials). The task was programmed and administered using Psychtoolbox version 3 (http://psychtoolbox.org/). The visual stimuli were projected on a screen located behind the scanner using PROPixx DLP LED projector (VPixx Technologies, Canada). The participants viewed the projected visual stimuli through a mirror attached to a head coil.

### fMRI procedure and preprocessing

MRI scanning was performed using a 3T MRI scanner (Siemens Prisma, Germany) with a 64-channel head coil. Functional images were acquired using an echo-planar imaging sequence [repetition time (TR): 2.0 s; echo time (TE): 27 ms; flip angle (FA): 90 deg; 36 slices; slice thickness: 3 mm; in-plane resolution: 2 × 2 mm]. Multiband accelerated imaging was unavailable because the guideline for the simultaneous fMRI–EEG recoding by the developer of the MRI-compatible EEG system did not permit it (Brain Products, Germany). Each functional run involved 265 volume acquisitions. The first five volumes were discarded for analysis to take into account the equilibrium of longitudinal magnetization. High-resolution anatomical images were acquired using an MP-RAGE T1-weighted sequence (TR: 2,400 ms; TE = 2.32 ms; FA: 8 deg; 192 slices; slice thickness: 0.9 mm; in-plane resolution: 0.93 × 0.93 mm).

All preprocessing was performed using SPM12 software (http://fil.ion.ac.uk/spm/). Functional images were firstly temporally realigned across volumes and runs, and the anatomical image was co-registered to a mean image of the functional images. The functional images were then spatially normalized to a standard Montreal Neurological Institute (MNI) template, with normalization parameters estimated for the anatomical scans. The images were resampled into 2-mm isotropic voxels. For the univariate fMRI analysis, functional volumes were spatially smoothed with a 6-mm full-width at half-maximum Gaussian kernel. For the deep learning analysis, we did not apply spatial smoothing to the fMRI data.

### EEG procedure and preprocessing

High-density EEG data were acquired from MRI-compatible 64 sintered Ag/AgCl electrodes with extended scalp coverage, including 63 scalp recording electrodes and an electrode on the upper back for electrocardiogram (ECG) (BrainCap-MR, Brain Products, Germany). Electrodes AFz and FCz served as the ground and online reference, respectively. All signals were recorded with a bandpass of 0.016–250 Hz and digitized at 5,000 Hz (BrainAmp MR Plus, Brain Products). The electrode impedances were lowered to <10 kΩ before recording and monitored throughout the experiment.

Offline preprocessing was firstly performed in Brain Vision Analyzer 2 (Brain Products). Initially, we removed the MR gradient artifact using an average template subtraction method (Allen et al., 2000) and down-sampled the resulting EEG data to 250 Hz. Secondly, the EEG data were bandpass filtered between 1 Hz and 100 Hz. Thirdly, the cardiovascular artifact was removed from the EEG data. Heartbeat detection using the ECG channel was then performed using a semiautomatic template-matching procedure. Fourthly, eye blink correction was performed using independent component analysis (ICA). As specific sensors for vertical/horizontal electrooculography (EOG) were not used, we employed the EEG data obtained from electrode FP1 as a substitute for the vertical activity of the EOG channel. Finally, we applied ICA-based denoising techniques implemented in EEGLAB (*runica* function). Then, each independent component was labeled using ICLabel plugin (https://github.com/sccn/ICLabel). Components with <75% probability of being in the brain signal were flagged as artifacts and rejected.

The resulting artifact-removed EEG data were further processed in EEGLAB as follows. If the signal was continuously noisy in a certain channel in a scanning run, the EEG data were interpolated at the corresponding channel by replacing it with the average EEG data from the neighboring channels. Accordingly, 16 of the 3,150 channels (50 participants × 63 scalp channels) were interpolated. Thereafter, the EEG data were segmented into 1.5 s epochs (0.5 s pre-stimulus to 1.0 s post-stimulus) time locked to the stimulus onset and were baseline-normalized. The data were then inspected manually to detect any trials containing any excessive noises. After excluding 742 trials, 17,265 trials were included in the analyses. The data were then digitally re-referenced to the average of all scalp channels.

For the ERPs, we firstly trial-averaged the EEG time series for each EEG channel separately. Then, the resulting ERPs were grand-averaged across participants to obtain population ERPs for each EEG channel. The spectral power was also calculated using the *mtspecgramc* function in the Chronux toolbox for MATLAB (http://chronux.org/). The parameters for the multi-taper time-frequency spectrum were as follows: window = 200 ms, step size = 80 ms, time-bandwidth product = 3, and number of tapers = 5.

### Respiration and eye-tracking recording

Respiratory data were recorded by using the BIOPAC MP160 system with AcqKnowledge software (BIOPAC Systems, Inc., Goleta, CA). Raw physiological time series of respiration were sampled at 2,000 Hz and then were preprocessed in the PhysIO toolbox (Kasper et al., 2017) for MATLAB as follows. Firstly, the respiration data were down-sampled to 250 Hz and bandpass-filtered between 0.1 Hz and 5.0 Hz. Then, both the respiratory phase and respiratory volume per time (RVT) for the respiratory data were obtained. From these physiological determinants with noise modeling, (1) 8 RETROICOR (RETROspective Image CORrection) regressors were computed via a fourth-order Fourier expansion of the respiratory phase (Harvey et al., 2008), and (2) respiratory response was calculated by convolution of the RVT with the respiration response function (Birn et al., 2006). Accordingly, we obtained a total of nine regressors and treated them as nuisance regressors in the fMRI univariate analysis (see “*fMRI univariate analysis*” section below).

To check whether the participants fixated their eye positions, the eye position during task runs was monitored online at a sampling rate of 1,000 Hz using an MRI-compatible infrared camera-based eye tracker (EyeLink 1000, SR Research, Ontario, Canada).

### DNN classifier

To explore spatiotemporal neural activity representing visual object category and sub-category, in-house DNN classifiers (Simonyan and Zisserman, 2015) were constructed (Figure 2). The architecture of the models is described below. The model training, validation, and testing framework for binary classification (face/object, male face/female face, or natural object/artificial object images) were implemented using PyTorch (https://pytorch.org/). The scripts employed for network modeling and analyses are available from GitHub (https://github.com/masaki-takeda/dld).

The whole dataset included 17,265 trial-based concurrent fMRI–EEG data from 50 participants. The data from 45 participants were used as a training/validation dataset for optimizing the hyperparameters, and independent data from five participants were used as a hold-out test dataset. In the training/validation dataset, data from five participants were treated as a unit of the fold, resulting in nine-fold datasets (Figure 2B). After preliminarily determining network architectures, we conducted grid searches for hyperparameter tuning by using the train/validation dataset (see “*Grid search*” section). After completing hyperparameter tuning, we next evaluated the generalization of neural decoding performance for the independent test dataset: nine-fold cross-validation was performed for the training/validation dataset, and then the resulting nine learned models were applied to the hold-out test dataset. For each binary classification for the test dataset, we equalized the number of trials for face/object, male face/female face, and natural object/artificial object by excluding a surplus of trials in the test dataset. This approach resulted in a chance level of 50% (categorization: 734 face trials vs. 734 object trials; face sub-categorization: 386 male face trials vs. 386 female face trials; object sub-categorization: 359 natural object trials vs. 359 artificial object trials). The decoding performance for visual object categorization and sub-categorization was then investigated using the following network models based on fMRI data alone (Figure 2C), EEG data alone (Figure 2D–E), and combined fMRI–EEG data (Figure 5A).

#### fMRI DNN

The current DNN model for fMRI data (fMRI DNN) consisted of a 3D convolutional neural network (CNN) (Frey *et al*., 2021) (Figure 2C). The 3D CNN included three convolution layers (*conv3d* function) for extracting spatial features and two fully connected layers. We also examined other networks with a greater number of convolution and fully connected layers, but the decoding performance did not change drastically. Based on the preliminary test for the training/validation dataset, the following hyperparameters and settings were selected: batch size of 100, early stopping, and Adam optimizer (Kingma and Ba, 2015). We then grid-searched for the number of averaged trials, learning rate, and weight decay.

The input datasets were trial-based non-smoothed fMRI volumes and were prepared as follows. To consider the delayed hemodynamic response of the BOLD signal, an fMRI volume that was acquired at 4 s after the onset of the visual stimulus presentation was collected (Figure 2A). The input fMRI volume was spatially normalized to the standard MNI space, and the signal intensities outside of the brain were masked to zero. Then, the signal intensity in each voxel was z-scored by the mean and SD of the signal intensity in the volume.

#### EEG DNN

Seven DNN models for EEG data (EEG DNNs) were tested in the preliminary inquiry for the training/validation dataset: (1) a custom DNN model including four *conv1d* layers, (2) a custom DNN model including four *conv2d* layers for band-passed EEG time series (delta, theta, alpha, beta, and gamma frequency band × time), (3) a recurrent neural network (RNN) model using long short term memory (LSTM), (4) a convolutional RNN (ConvRNN) model using LSTM after two *conv1d* layers, (5) a spatial-temporal neural network originally designed for P300 detection (Zhang et al., 2021), (6) a generic temporal convolutional network (TCN) (Bai *et al*., 2018), and (7) a modified version of the TCN. Among them, we adopted the TCN model (6) because of its superior decoding performance and its simplicity (Figure 2D– E). The TCN model has several distinguishing characteristics, such as dilated convolutions (dilated *conv1d*) and the causal architecture (no information leakage from future to past). Based on the preliminary test, a batch size of 100 and early stopping were selected. We then grid-searched for the number of averaged trials, learning rate, and kernel size. The dilation factors for each kernel size were determined to cover all the input time series.

The input datasets were trial-based EEG time series data and were prepared as follows. Preprocessed EEG time series data were collected from 63 scalp channels. The segmented data obtained at 1 s from the stimulus onset (250 samples at a sampling rate of 250 Hz) were used (Figure 2A). The resulting segmented data were further z-scored by the mean and SD of the signal amplitude for each channel.

#### DNN for combined data

We also tested whether concurrent usage of both fMRI and EEG data improves the decoding performance relative to the usage of fMRI or EEG data alone. The DNN architecture for combined data (fMRI–EEG DNN) is shown in Figure 5A. In the model, the feature extraction layers of the two networks for fMRI and EEG data were concatenated and then fully connected. The fMRI and EEG networks were pre-trained separately, and the initial parameters of the convolution layers in the fMRI–EEG DNN were set to the pre-trained parameters.

#### Grid search

Before conducting the grid search, we preliminarily searched several hyperparameters such as the number of layers and kernel sizes, and found that several hyperparameters affected the decoding performance: the number of averaged trials, weight decay, and learning rate for the fMRI data, and the number of averaged trials, kernel size for delated convolutional layers, and learning rate for the EEG data. Thus, grid searches were conducted to obtain the optimal combination of these hyperparameters. For the fMRI DNN, the following hyperparameters were tuned: the number of averaged trials (1, 3, 5, 7, or 9), learning rate (0.0001–0.01 in five divisions), and weight decay (0–0.01 in five divisions). For the EEG DNN, the following hyperparameters were tuned: the number of averaged trials (1, 3, 5, 7, or 9), learning rate (0.0001–0.01 in 5 divisions), and kernel size (2–9 in five divisions). For the combined fMRI–EEG DNN, the following hyperparameters were tuned: the number of averaged trials (1, 3, 5, 7, or 9), learning rate of feature extraction layers for fMRI data (0.001 or 0.0001), weight decay for fMRI data (0 or 0.001), learning rate of feature extraction layers for EEG data (0.0001 or 0.0005), and kernel size for EEG data (2, 5, 7, or 9). For the combined fMRI–EEG DNN, we fixed the learning rate and weight decay for the fully connected layers as 0.001 and 0, respectively. All grid search analyses were conducted using the training/validation dataset, and the hold-out test dataset was completely free from the hyperparameter settings.

#### Data averaging

A single-trial neural signal contains noise derived from the brain and measurement hardware, and one of the methods of increasing the signal to noise (S/N) ratio is to average the data. To estimate how the S/N ratio of brain signals affects the decoding performance, we tested the decoding using both data from a single trial data and averaged data (number of averaged trials ≤9). We averaged the fMRI or EEG data within each subject and then augmented the data to match the total average number of data with the number of all trials (e.g., averaged data from nine trials were augmented by a factor of nine).

#### Visualization of relevant neural activity in DNNs

To highlight important spatiotemporal neural activities for visual object classification, visualization approaches in deep learning were applied. For both the fMRI DNN and EEG DNN, guided Grad-CAM (Selvaraju *et al*., 2019) was used to create a high-resolution class-discriminative visualization. Guided Grad-CAM was created by multiplying guided backpropagation at the input of the first convolution layer with Grad-CAM at the last convolution layer. The original Grad-CAM uses global average pooling of the gradients (Selvaraju *et al*., 2019), but we did not use it like Layer-CAM (Jiang et al., 2021), because the smoothing effect of pooling was extremely strong, especially for the 3D convolution. In the end, we obtained multiple guided Grad-CAM maps, the number of which corresponded to the number of trials. The obtained guided Grad-CAM values were not normalized. As for EEG guided Grad-CAM, the model without residual connections was used because those connections can cause gridding artifacts with the dilated convolution layers and obfuscate the visualization results (Yu et al., 2017). The resulting guided Grad-CAM values in the EEG DNN were then smoothed using the Gaussian-weighted moving average with a 40-ms window.

### Statistics

#### General

All statistical analyses were performed in MATLAB and Python. All data are shown as mean ± standard error of the mean unless otherwise stated. The sample sizes were not based on a priori power calculations but are comparable to those employed in other studies in the field using similar techniques (Supratak *et al*., 2017).

#### fMRI univariate analysis

The fMRI univariate analysis was conducted as in previous studies (Matsui et al., 2022; Tanaka et al., 2020; Tsumura et al., 2021a; Tsumura *et al*., 2021b) (Figure 3A–C). In the first-level analysis, a GLM (Worsley and Friston, 1995) approach was used to estimate parameter values for task events. The events of interest were the presentation of male face, female face, natural object, artificial object stimuli (four regressors). Only correct trials were coded as regressors. Those task events were time locked to the onset of stimulus presentation and then convolved with a canonical hemodynamic response function implemented in SPM. Additionally, the onset of blank stimuli, six-axis head movement parameters, white matter signal, CSF signal, and effect of respiration (nine regressors) were included in GLM as nuisance effects, totaling to 18 nuisance regressors. Then, parameters were estimated for each voxel across the whole brain.

In the group level analysis, maps of parameter estimates were first contrasted within individual participants. The contrasted maps were then collected from participants and subjected to group-mean tests. Voxel clusters were first identified using a voxel-wise uncorrected threshold of *P* < 0.001. The voxel clusters were then tested for significance across the whole brain with a threshold of *P* < 0.05 corrected by the family-wise error (FWE) rate based on permutation methods (5,000 permutations) implemented in the *randomise* function in the FSL suite (http://fmrib.ox.ac.uk/fsl/), which was empirically validated to appropriately control the false positive rate in a previous study (Eklund et al., 2016). Peaks of significant clusters were then identified and listed in tables. If multiple peaks were identified within 12 mm, the most significant peak was retained.

#### EEG univariate analysis

For each participant, a single ERP and power spectrogram were constructed by averaging the data across all trials separately for each visual category or sub-category in each channel. We then averaged the ERPs/spectrograms from all participants, which resulted in a grand average of 63-ch ERPs/spectrograms for the face, object, male face, female face, natural object, and artificial object trials (Figure 4A–C; Figure 4 – figure supplement 1A). The subsequently obtained spectrograms were further bandpass-filtered to divide them into alpha (8–15 Hz), beta (15–30 Hz), and gamma (30–70 Hz) frequency bands.

For statistical inference of differential EEG time series of ERP and power spectrograms (e.g., face minus object trials) against zero, permutation-based cluster-size inference (Dobs *et al*., 2019; Maris and Oostenveld, 2007) (i.e., a cluster refers to a set of contiguous time points) was performed separately for each channel. Time points that exceeded the 97.5^th^ percentile of the permutation distribution (*N* = 5,000) served as cluster-inducing time points (i.e., equivalent to *P* < 0.05, two-sided). The significance of clusters was set at the threshold of *P* < 0.05 (Figure 4 – figure supplement 1B–D).

#### Neural decoding performance

We trained our DNN models using nine-fold cross-validation with the training/validation dataset and then evaluated their generalization performance with the test dataset. Thus, a total of nine generalization performances were obtained for the fMRI DNN model, EEG DNN model, and combined fMRI–EEG DNN model. Three different statistical tests were performed to examine the significance of decoding accuracies and their robustness (Figure 3D, 4D, and 5B). Firstly, an ensemble learning method with majority voting (Rokach, 2009) was conducted. In this scheme, each base model (*N* = 9) made a prediction and voted for each sample (a stimulus in a trial). Only the sample class with the highest votes was included in the final predictive class. Statistical significance was tested using the binomial test against chance level (50%). Secondly, the t-test was conducted to determine the distribution of accuracies obtained by the nine models against the chance level, with FDR corrections (*q* < 0.05). Thirdly, the binomial test was performed for each model against chance level, with FDR corrections (*q* < 0.05), and the number of significant models was counted.

#### Visualization of neural decoding

The model that showed the maximum accuracy in the average of nine trials was selected for the subsequent visualization analysis.

The statistical significance of the guided Grad-CAM for fMRI data was evaluated by conducting the permutation-based cluster-size inference (Dobs *et al*., 2019; Maris and Oostenveld, 2007) used in the present univariate analyses (Figure 3E; see “*fMRI univariate analysis*” section). The spatial distribution of the significant guided Grad-CAM was also evaluated based on the Harvard-Oxford cortical and subcortical structural atlases, which covered 48 cortical and 21 subcortical structural areas, implemented in FSL (Figure 3 – figure supplement 4; *HarvardOxford-cort-maxprob-thr50-2mm* and *HarvardOxford-sub-maxprob-thr50-2mm*). We divided the cortical areas into areas in the right and left hemispheres and obtained 95 cortical regions (note that there were no voxels for the left supracalcarine cortex). Regarding the subcortical areas, we bilaterally removed areas labeled as the cerebral white matter, lateral ventricle and cerebral cortex and obtained 15 subcortical regions. We examined whether the guided Grad-CAM values in each brain regions (*N* = 110) were significantly higher than those obtained when DNNs were not learned. In the non-learned model, the weight values were initialized and fixed as random values. Statistical significance was evaluated by performing the Wilcoxon signed rank test with FDR corrections (*q* < 0.05).

We also conducted two types of correlation analysis (Spearman’s rank correlation; Figure 3 – figure supplement 5) and three types of ROI analysis to examine the spatial overlap between activations in the univariate analysis and guided Grad-CAM (Figure 3F, Figure 3 – figure supplement 6). The correlation was calculated either for the voxels masked by the gray matter prior volume of the FSL (*avg152T1_gray* template, threshold = 0.5) or for the voxels masked by the maps of meta-analysis obtained from Neurosynth (Yarkoni *et al*., 2011) (https://neurosynth.org) using the terms “face” and “object” (uniformity test). In the ROI analysis, we examined whether the guided Grad-CAM values in each ROI were significantly higher than those obtained using the non-learned DNNs. The ROIs were selected based on (1) the univariate analysis (Figure 3 – figure supplement 1; Figure 3F), (2) the anatomical structures from the Harvard–Oxford atlas (Figure 3 – figure supplement 6A), or (3) the maps of meta-analysis obtained from Neurosynth (Figure 3 – figure supplement 6B). In the Harvard-Oxford atlas (2), we created ROIs that included the peak voxels of the clusters identified by the univariate analysis, by thresholding the probabilistic map of the region by 50%. Note that we combined MFGi and MFGs for object trials identified in the univariate analysis into one ROI as MFG. In the Neurosynth (3), ROIs were selected with the uniformity test using the terms “face” and “object.” ROIs with more than 50 voxels were used in the analysis. In all ROI analyses, statistical significance was evaluated using Wilcoxon signed rank test with FDR corrections (*q* < 0.05).

The guided Grad-CAM for EEG data were quantified according to two indices, emergence latency and maximum latency. Firstly, the guided Grad-CAM values for each trial were averaged across 63 channels. Then, for each trial type, we defined the emergence latency as the time point at which the value first exceeded the level at 20% of the maximum of channel-averaged guided Grad-CAM values (Figure 4E). We obtained similar results when the threshold was changed to 10%, 15%, 25%, and 30%. The maximum latency for each trial was also defined as the time point at which the guided Grad-CAM showed maximum value in each trial (Figure 4F), and statistical significance in the maximum latency across categorization/sub-categorization was tested using the Kruskal–Wallis test with the post-hoc Tukey-Kramer test (*P* < 0.05). We also calculated the maximal latencies and corresponding guided Grad-CAM values when the DNN was not learned. Then, we examined whether the guided Grad-CAM values at the maximum latency were significantly higher in the learned model than in the non-learned model using the Wilcoxon signed rank test with FDR corrections (*q* < 0.05; Figure 4G). Finally, we tested whether the guided Grad-CAM values at the maximum latency differed across brain regions. EEG channels were grouped into six brain regions (frontal cortex, central cortex, parietal cortex, occipital cortex, left temporal cortex, and right temporal cortex; Figure 4 – figure supplement 2). The electrodes for the six regions are as follows: Fp1, Fp2, Fz, F1, F2, AF3, AF4, AF7, AF8, and Fpz for the frontal region; C3, C4, Cz, FC1, FC2, C1, C2, FC3, and FC4 for the central region; P3, P4, Pz, CP1, CP2, P1, P2, CP3, CP4, P5, P6, and CPz for the parietal region; F3, F7, T7, P7, FC5, CP5, TP9, F5, C5, FT7, TP7, and FT9 for the left temporal region; and F4, F8, T8, P8, FC6, CP6, TP10, F6, C6, FT8, TP8, and FT10 for the right temporal region. The values obtained by each channel in a region were averaged, and regional differences were tested using Friedman’s test with the post-hoc Tukey–Kramer method (*P* < 0.05).

### Data and code availability

All data supporting the findings of this study are provided within the paper. All custom Python codes used for deep learning and learned model data in this study are available at https://github.com/masaki-takeda/dld. All additional information will be made available upon reasonable request to the authors.

## Acknowledgments

This study was supported by Kakenhi (Japan Society for the Promotion of Science) grant numbers 20H00521 and 21K18267 to MT; 21H00211 and 21K07262 to NW; 21H05060 to KJ; 21K20303 to DS; and 17H00891 to KN. This study was also supported by a grant from Uehara Memorial Foundation to MT, and a grant from Takeda Science Foundation to MT. We thank Dr. Shigeyuki Oba, Dr. Shinichi Yoshida, Mr. Zhen Zhang, and Dr. Makoto Iwata for their technical advice on deep learning. We also thank Mr. Rei Furutani and Ms. Maoko Yamanaka for their technical and administrative assistance, respectively.

## Author contributions

N.W., K.J., K.N., and M.T. designed the experiment and study. N.W., R.K., and M.T. collected the data. N.W., K.M., and M.T. analyzed the data. N.W., K.M., K.J., D.S., K.N., and M.T. wrote the manuscript.

---

**Figure 1 – figure supplement 1.**
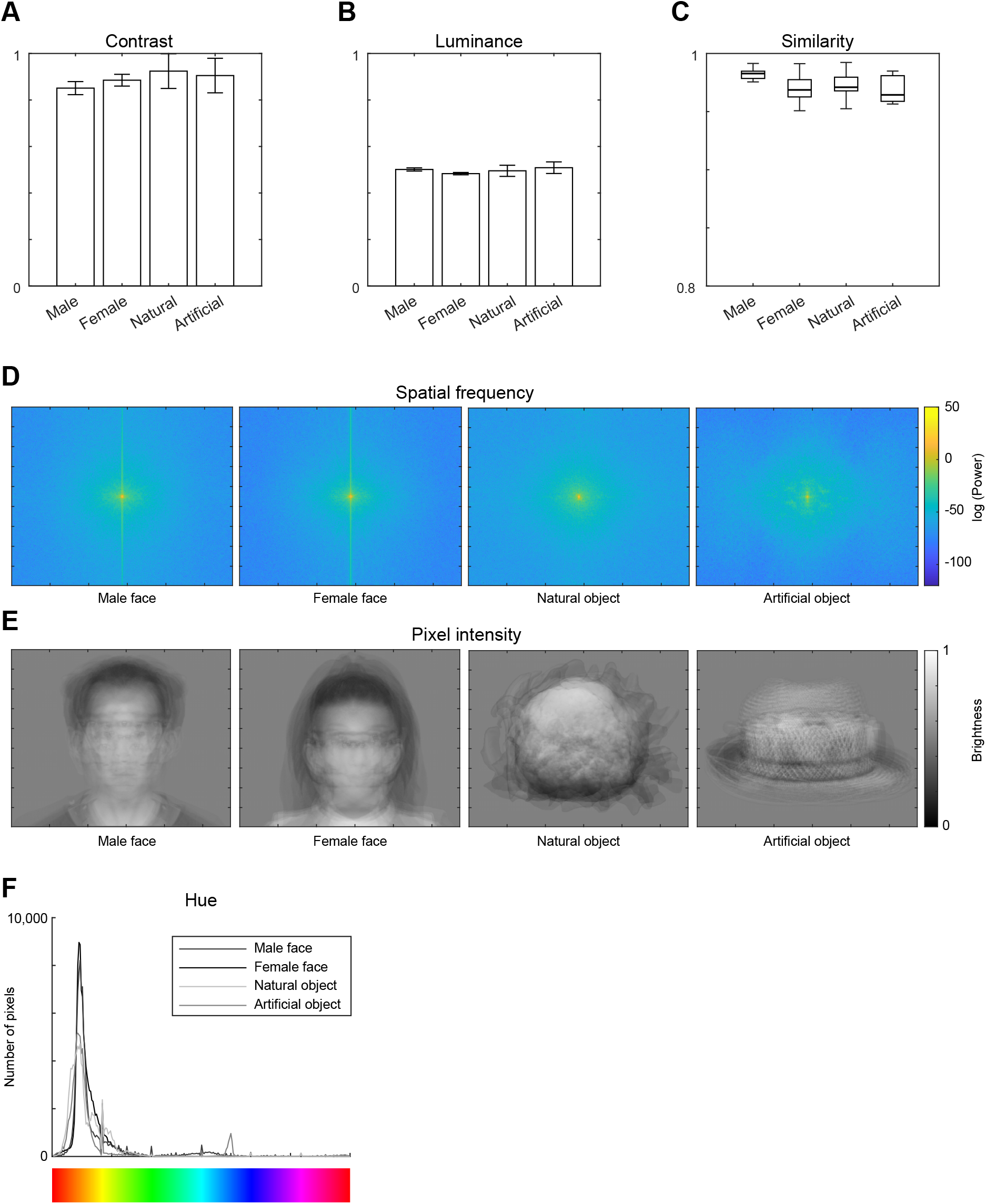
Low-level visual features of stimulus images. **(A)** Michelson contrast of images as the ratio between the difference and the sum of the maximum and minimum pixel intensities. **(B)** Luminance as the mean pixel intensity of images. **(C)** Pixel-based pairwise similarity. To estimate the visual similarity among the images in a category, the mean Euclidean distance between normalized grayscale values (scaled to be between 0 and 1) of each stimulus and every other stimulus of the same sub-category was measured. The pairwise similarity values range from 0 (images with inverted intensity difference at each pixel) to 1 (identical images). Note that the order of the size of the effect [0.0062, standard deviation (S.D.) of the category means in normalized grayscale values] is comparable to the resolution of the display (0.0039, minimum pixel-wise distance resolved by the display in normalized values, corresponding to the change in one gray-level value of a possible 255 increments). Mean ± S.D. are shown in (A–C). **(D)** Average spatial frequency across images. **(E)** Average pixel intensity for each pixel. **(F)** Average hue histogram across images.

**Figure 3 – figure supplement 1.**
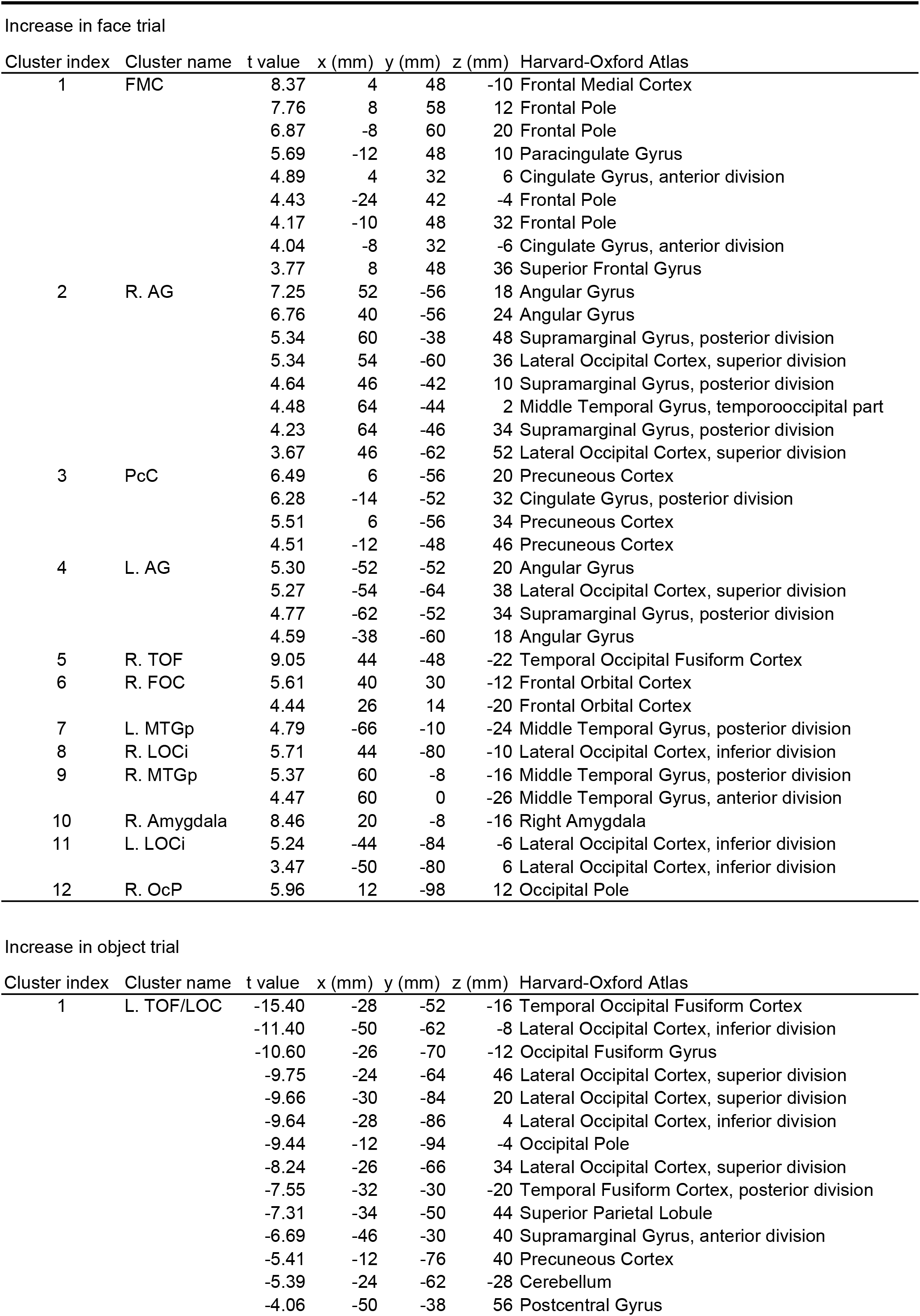
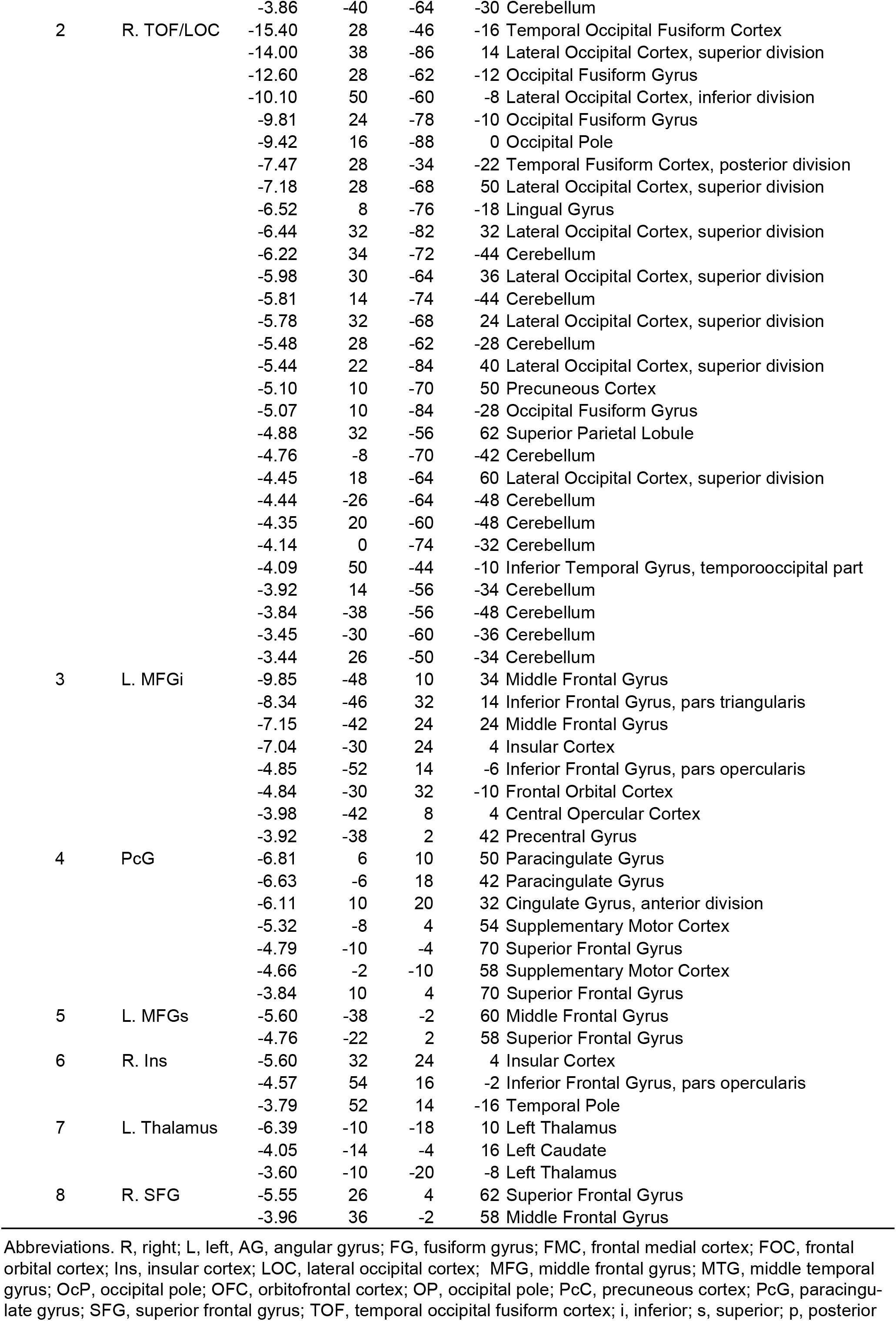
Brain regions showing significant signal increases and decreases in the contrast of face vs. object. Coordinates are listed in MNI space.

**Figure 3 – figure supplement 2.**
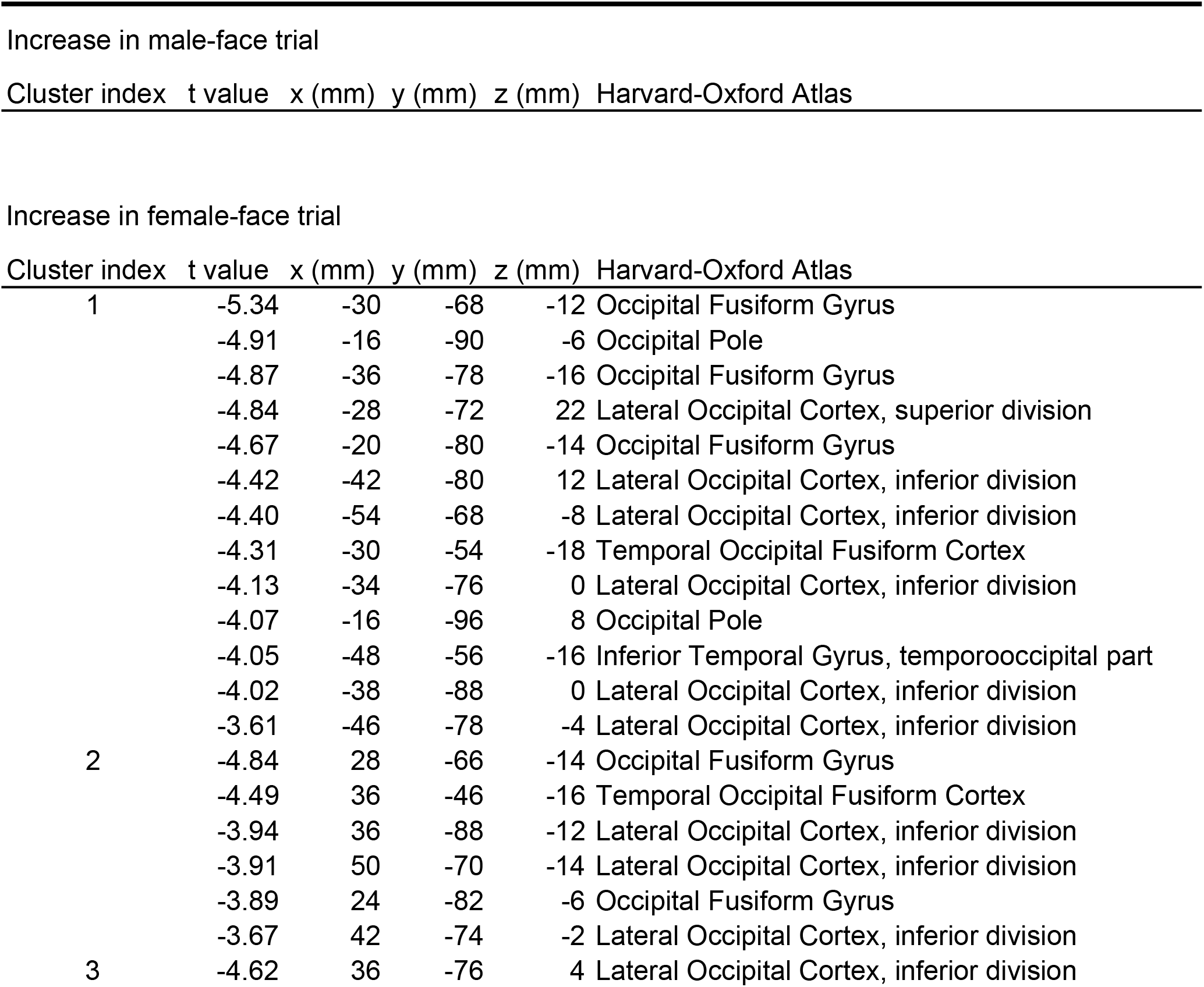
Brain regions showing significant signal increases and decreases in the contrast of male-face vs. female-face. Coordinates are listed in MNI space.ase in male-face trial

**Figure 3 – figure supplement 3.**
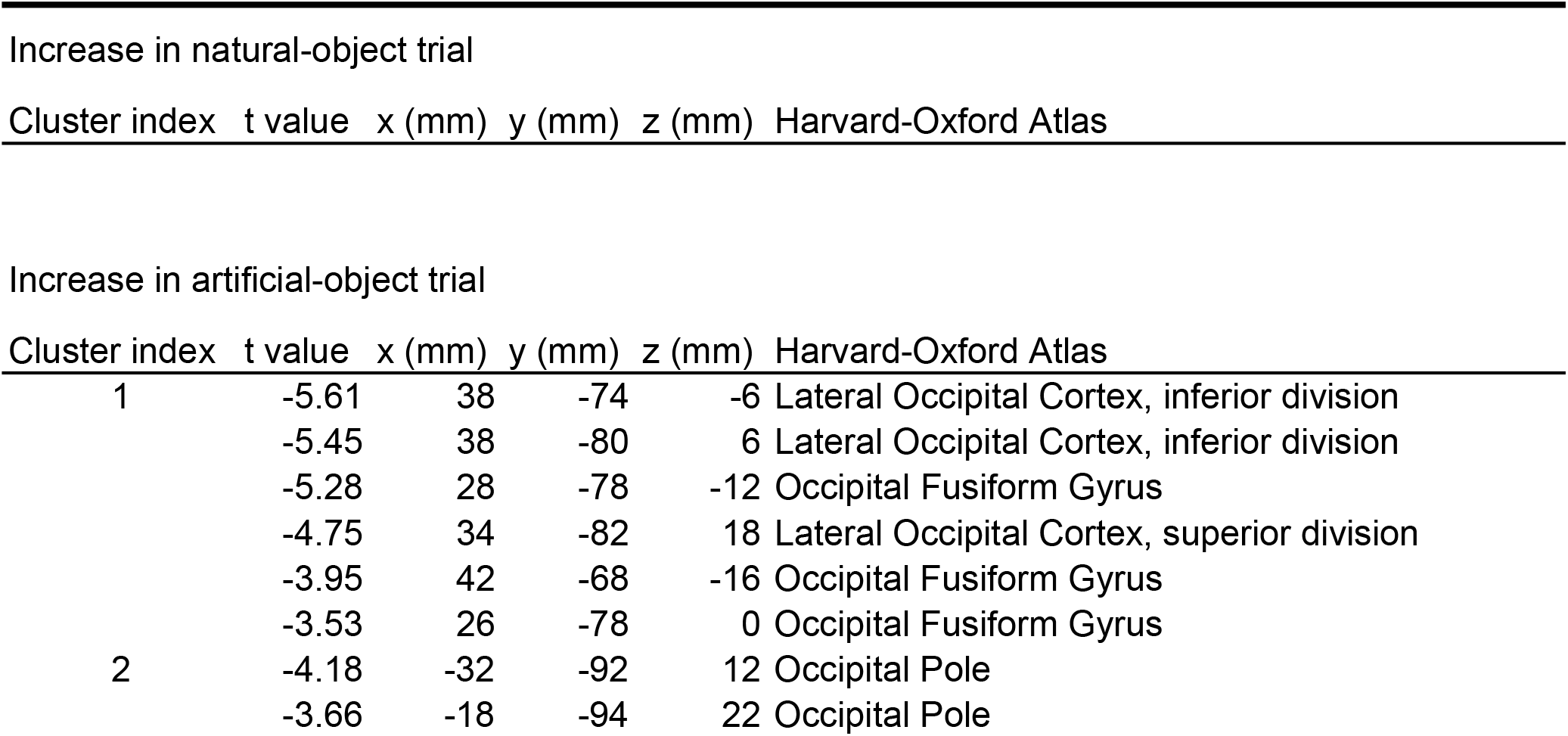
Brain regions showing significant signal increases and decreases in the contrast of natural-object vs. artificial-object. Coordinates are listed in MNI space.

**Figure 3 – figure supplement 4.**
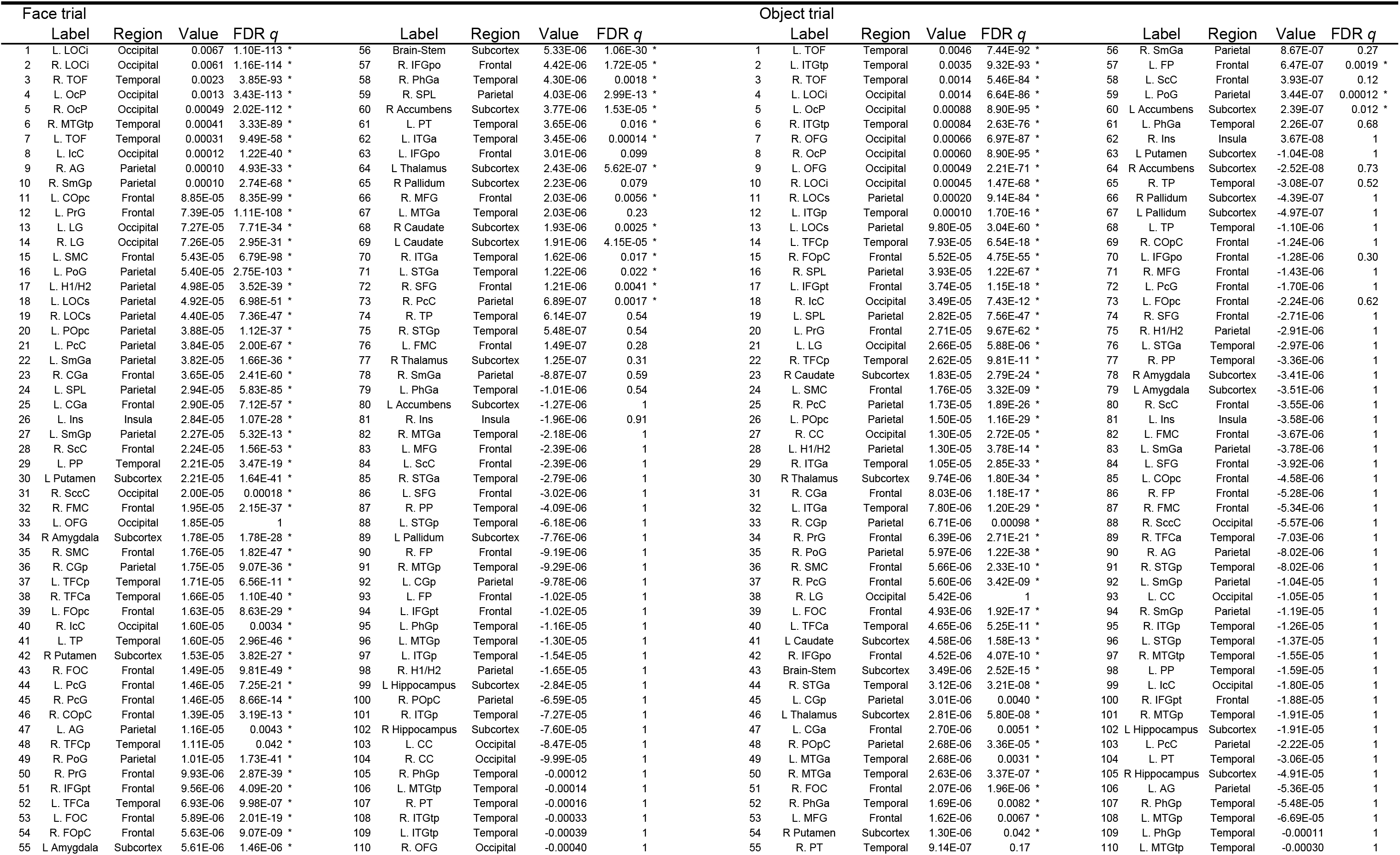
Guided Grad-CAM values for face and object trials in anatomically-defined brain regions. The brain regions are sorted by the magnitudes of the values. An asterisk denotes FDR q < 0.05.

**Figure 3 – figure supplement 5.**
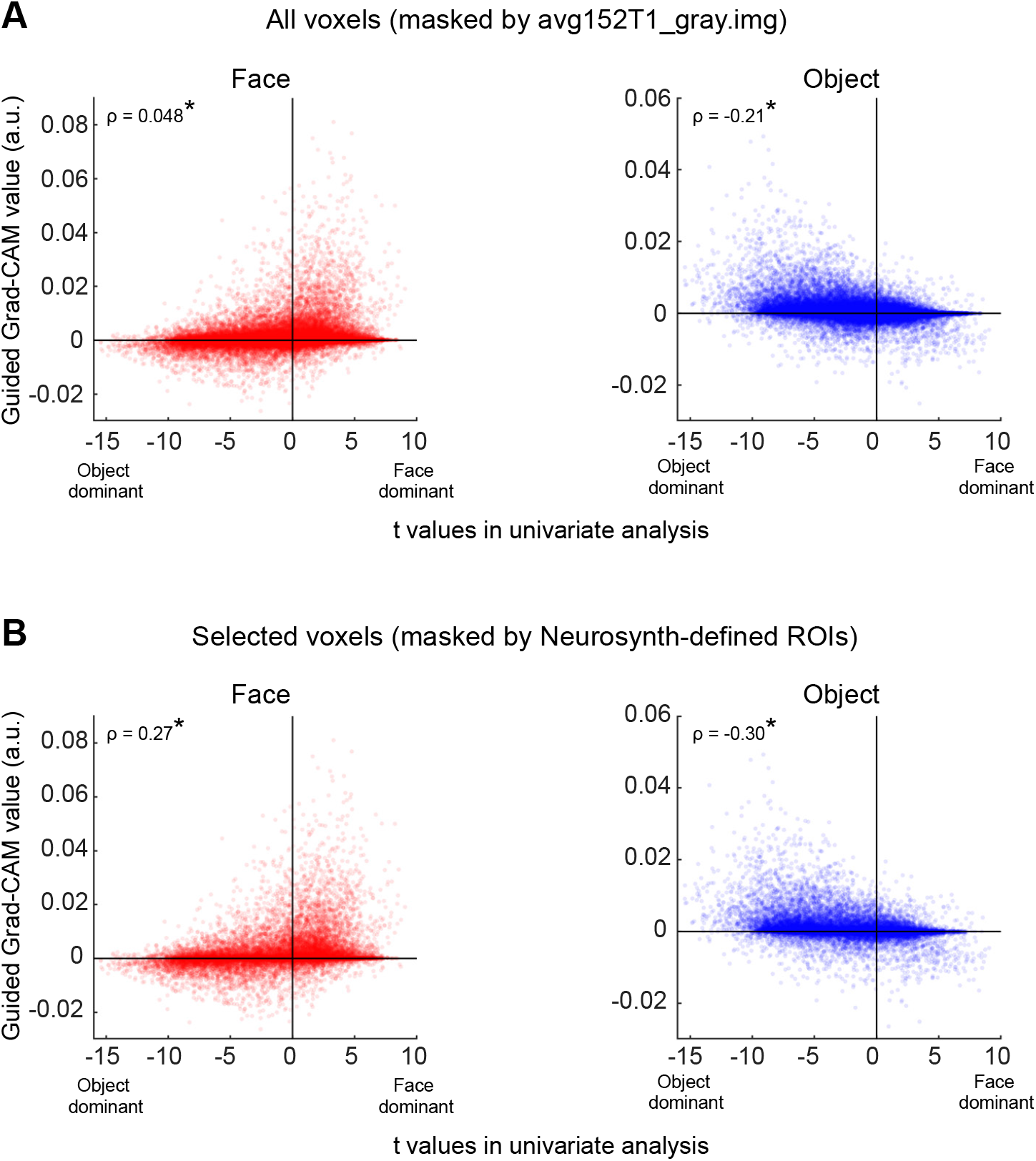
Voxel-based relationship between t-values in the univariate analysis and the guided Grad-CAM values in the face and object trial. **(A)** Relationship between the two values when voxels are masked by the gray matter prior volume of the FSL. **(B)** Relationship between the two values when the voxels are masked by functional ROIs obtained from Neurosynth using the terms “face” and “object.” *, *P* < 0.001.

**Figure 3 — figure supplement 6.**
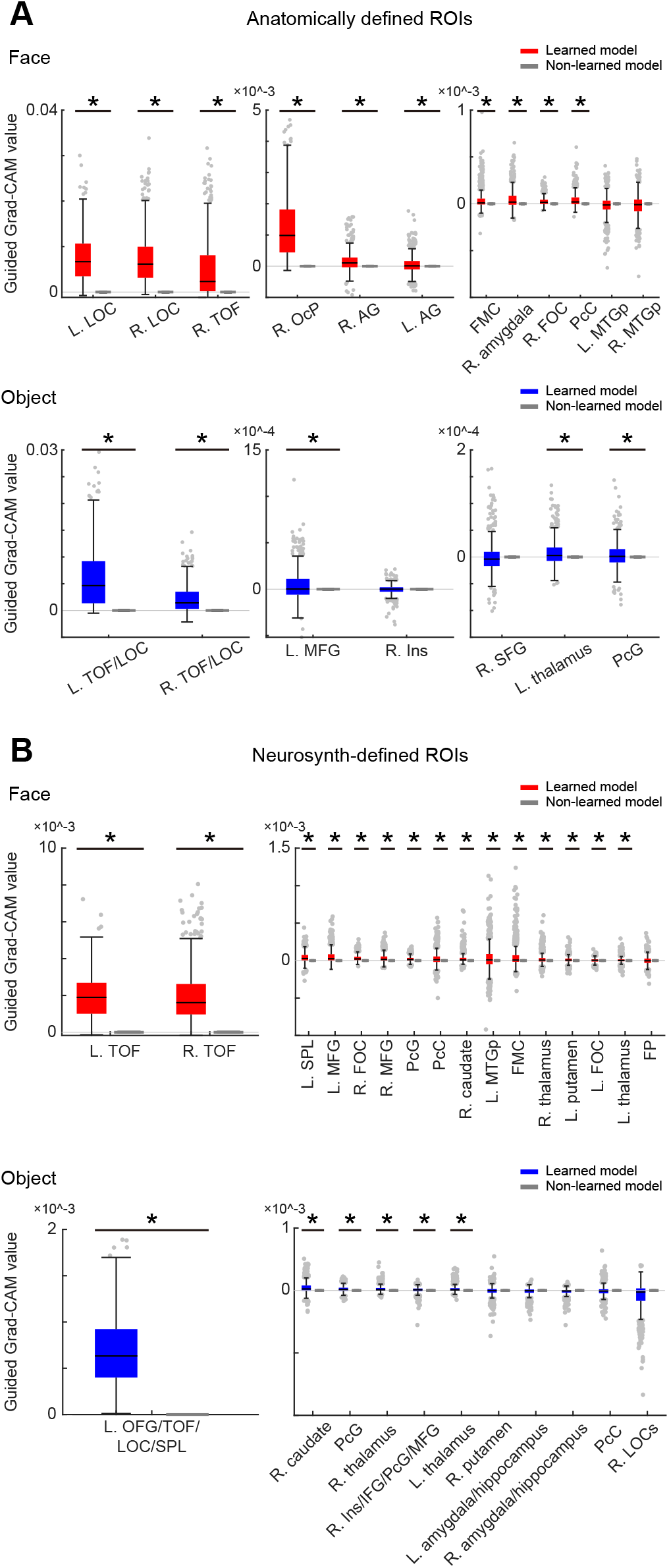
ROI analysis. Guided Grad-CAM values for each ROI when ROIs are defined **(A)** anatomically (Harvard–Oxford atlas) and **(B)** functionally (Neurosynth). The figure configurations are the same as those in Figure 3f. L, left; R, right; AG, angular gyrus; FMC, frontal medial cortex; FOC, frontal orbital cortex; FP, frontal pole; IFG, inferior frontal gyrus; Ins, insular cortex; LOC, lateral occipital cortex; MFG, middle frontal gyrus; MTG, middle temporal gyrus; OcP, occipital pole; OFG, occipital fusiform gyrus; PcC, precuneous cortex; PcG, paracingulate gyrus; SFG, superior frontal gyrus; SPL, superior parietal lobule; TOF, temporal occipital fusiform cortex; i, inferior; s, superior; p, posterior.

**Figure 4 — figure supplement 1.**
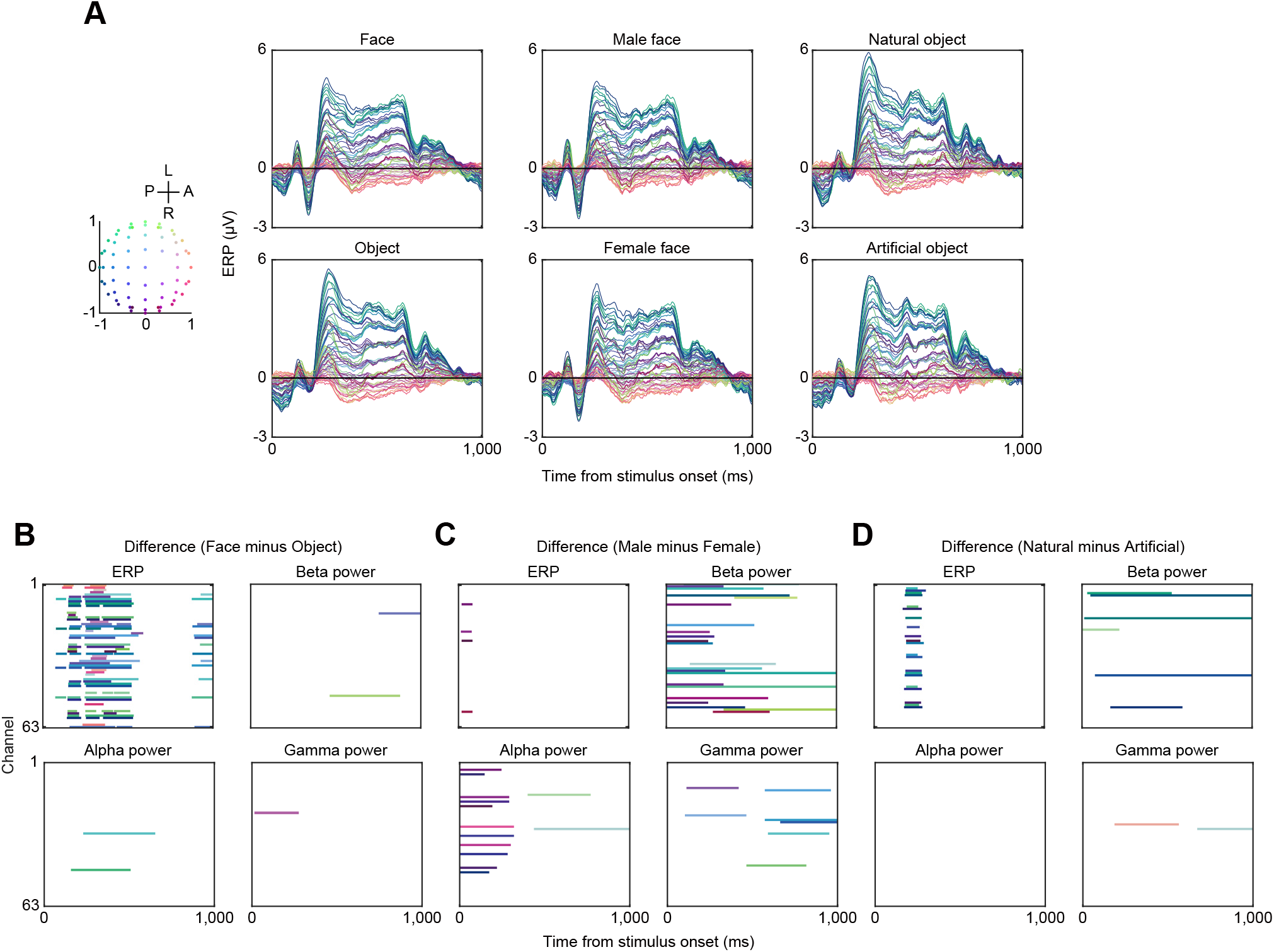
EEG responses. **(A)** ERPs for face trials, object trials, male face trials, female face trials, natural object trials, and artificial object trials. The line colors correspond to the channel locations depicted on the left side. A, anterior, P, posterior, R, right, L, left. **(B–D)** Significance of temporal trajectories in ERPs and power spectrograms. The colored lines depict the time points showing significant differential ERP amplitudes or power spectrograms against zero in the respective channels. The color of each line depicts the location of the corresponding channel (left). Time points that exceeded the 97.5^th^ percentile of the permutation distribution (*N* = 5,000) served as cluster-inducing time points, and the significance of clusters was set at the threshold of *P* < 0.05.

**Figure 4 — figure supplement 2.**
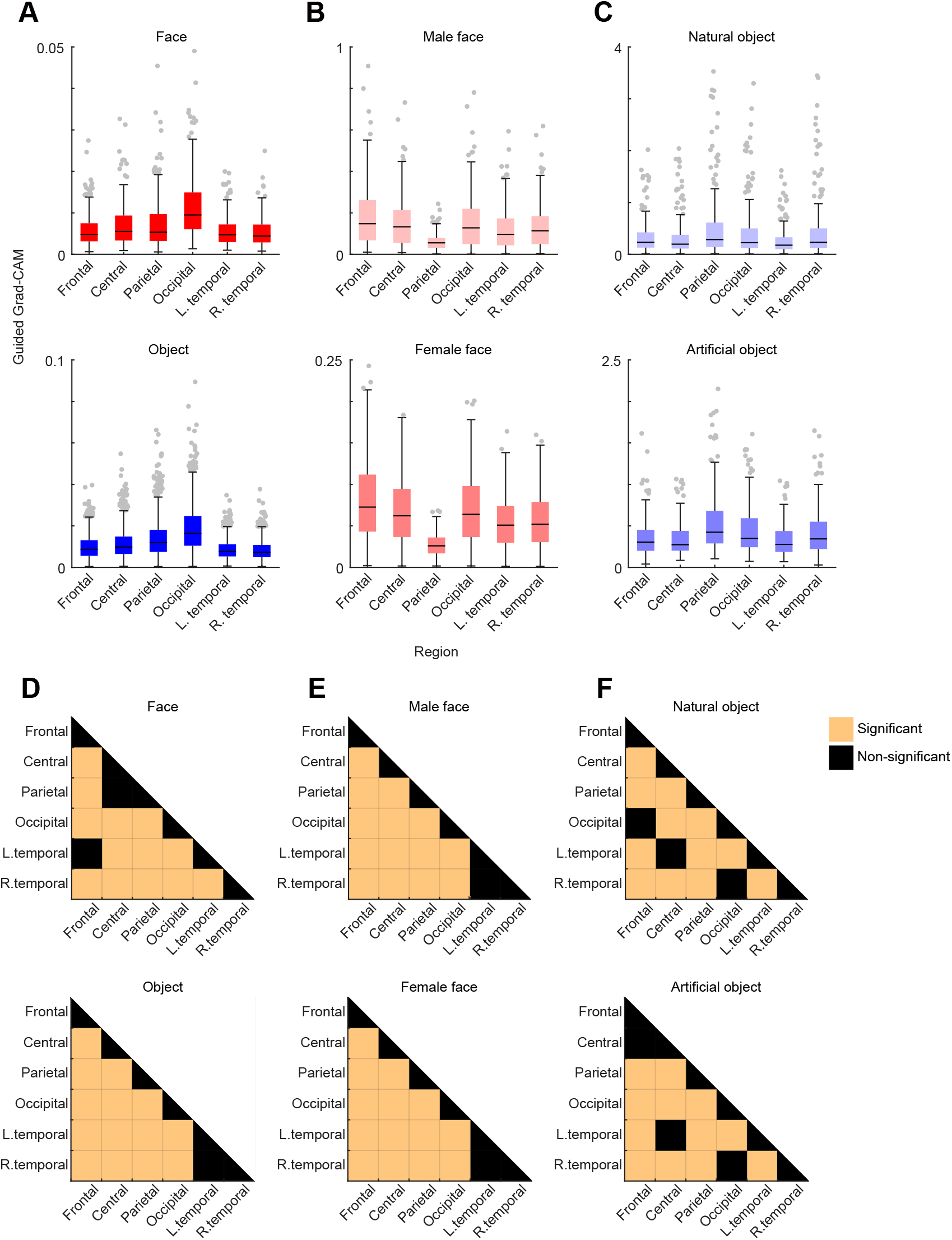
Regional analysis of guided Grad-CAM. **(A–C)** Comparison of the guided Grad-CAM values at the maximum latency in face and object trials (A), male face and female face trials (B), and natural object and artificial object trials (C) across six regions. All 63 channels are categorized as either frontal, central, parietal, occipital, left temporal, and right temporal regions based on their locations. The trial-based median value distributions across channels in a region are shown. The regional differences in the maximum guided Grad-CAM values are statistically significant in all trial types (Friedman’s test, *P* < 0.001). **(D–F)** Results of multiple comparison test (Tukey–Kramer method, *P* < 0.05). Significant differences in the guided Grad-CAM values between regions are color-coded.

